# Repurposing the *Streptococcus mutans* CRISPR-Cas9 System to Understand Essential Gene Function

**DOI:** 10.1101/791483

**Authors:** RC Shields, AR Walker, N Maricic, B Chakraborty, SAM Underhill, RA Burne

## Abstract

A recent genome-wide screen identified ∼300 essential or growth-supporting genes in the dental caries pathogen *Streptococcus mutans*. To be able to study these genes, we built a CRISPR interference tool around the Cas9 nuclease (Cas9_Smu_) encoded in the *S. mutans* UA159 genome. Using a xylose-inducible dead Cas9_Smu_ with a constitutively active single-guide RNA (sgRNA), we observed titratable repression of GFP fluorescence that compared favorably to that of *Streptococcus pyogenes* dCas9 (Cas9_Spy_). We then investigated sgRNA specificity and proto-spacer adjacent motif (PAM) requirements. Interference by sgRNAs did not occur with double or triple base-pair mutations, or if single base-pair mutations were in the 3’ end of the sgRNA. Bioinformatic analysis of >450 *S. mutans* genomes allied with *in vivo* assays revealed a similar PAM recognition sequence as the Cas9_Spy_. Next, we created a comprehensive library of sgRNA plasmids that were directed at essential and growth-supporting genes. We discovered growth defects for 77% of the CRISPRi strains expressing sgRNAs. Phenotypes of CRISPRi strains, across several biological pathways, were assessed using fluorescence microscopy. A variety of cell structure anomalies were observed, including segregational instability of the chromosome, enlarged cells, and ovococci-to-rod shape transitions. CRISPRi was also employed to observe how silencing of cell wall glycopolysaccharide biosynthesis (rhamnose-glucose polysaccharide, RGP) affected both cell division and pathogenesis in a wax worm model. The CRISPRi tool and sgRNA library are valuable resources for characterizing essential genes in *S. mutans*, some of which could prove to be promising therapeutic targets.

## Introduction

*Streptococcus* species are natural inhabitants of humans and animals. Under the right conditions many streptococci can cause or exacerbate diverse infections in humans, including life-threatening illnesses that include meningitis or pneumonia [1]. The transition between commensalism and pathogenicity is an important feature of the oral streptococci, which is the most abundant bacterial genus in the human mouth [2]. Oral streptococci have important roles in the initiation and maturation of dental biofilms (plaque), and many are associated with oral and dental health. However, oral streptococci that are especially acidogenic and acid-tolerant are implicated as primary etiologic agents in the pathogenesis of dental caries (tooth decay) [3]. Dental caries is a common cause of pain, impaired quality of life, and tooth loss in children and adults [3]. The organism most often linked to caries development is *Streptococcus mutans*, which has been shown to have multiple virulence traits, some unique, that imbue the organism with the capacity to initiate and worsen the disease [4]. Extraorally, *S. mutans* and other oral streptococci can cause infective endocarditis [5]. More recently, epidemiologic and mechanistic studies point to a role for certain *S. mutans* strains in cardiovascular diseases, including stroke [6]. Investigating the biology of this organism is important for developing therapies that combat dental caries and other diseases associated with *S. mutans* and related organisms. More broadly, expanding the understanding of streptococcal biology will positively impact efforts to prevent or treat many other diseases in humans and animals.

Antimicrobial therapies usually target biological processes that are essential for bacterial growth and persistence. These pathways include protein synthesis, transcription, DNA replication, cell division, peptidoglycan biosynthesis, or folic acid metabolism. The most traditional approach for discovering new antibiotics has been screening the ability of natural and synthetic compounds to inhibit growth or processes essential for pathogenesis [7]. Alternatively, antibacterial targets could be identified by combining genome sequence data with genome-wide techniques that probe gene essentiality, followed by the identification or design of molecules that target the factors involved in the process of interest [8]. Global identification of essential genes is possible with a technique known as transposon sequencing (Tn-seq) [9]. Using Tn-seq, we identified a panel of genes that are essential for the growth of *S. mutans* UA159, together with genes that are required for growth *in vitro* in rich or defined medium [10]. Although Tn-seq can be used to discover essential genes, it does not yield conditional mutations, and therefore does not allow for the functional study of these genes unless culture conditions can be found that allow for growth. Before moving toward studies that attempt to block or inhibit these pathways in *S. mutans*, or other streptococci, there is a need for further validation and investigation of these apparently essential genes. However, functional studies of essential genes in bacteria have traditionally been technically challenging and time-consuming, because mutants of these genes cannot be isolated and studied. One approach involves the generation of a conditional knockout, usually by mutating an essential gene and simultaneously expressing a copy of the gene under the control of a promoter that can be activated by an exogenously provided compound [11]. Two major drawbacks to this approach are that it is not easily adapted to a comprehensive study of the essential genome of an organism in a time-efficient manner (low throughput) and differences in gene expression levels between the conditional expresser and the wild type can generate misleading results.

Reprogramming of the bacterial CRISPR (<u>c</u>lustered regularly interspaced <u>s</u>hort <u>p</u>alindromic repeats) -Cas system into a gene-silencing tool has made it more practical to perform high-throughput studies of essential genes [12,13]. This technique, known as CRISPR interference (CRISPRi), combines an inducible, catalytically inactive *Streptococcus pyogenes* Cas9 nuclease, dead Cas9 (dCas9), with a programmable single-guide RNA (sgRNA). The sgRNA encodes a 20-nt sequence that is complementary to the target DNA. When expressed together in a cell, the sgRNA-dCas9_Spy_ complex represses transcription initiation or elongation of the target gene. CRISPRi libraries that silence all putatively essential genes have been generated for both *Streptococcus pneumoniae* D39 [14] and *Bacillus subtilis* 168 [15]. A complicating factor for developing a CRISPRi system in *S. mutans* UA159 is the presence of an endogenous Cas9 nuclease (SMu.1405c) [16]. Introduction of a CRISPRi system into *S. mutans* built around dCas9_Spy_ would likely not function without the deletion of the endogenous Cas9, because of the potential for Cas9_Smu_-sgRNA complexes to cause lethal double-strand breaks in the chromosome. It is known that Cas9_Smu_ and Cas9_Spy_ can cleave DNA in the presence of a *S. pyogenes* dual-RNA, and vice versa [17]. The Cas9 target recognition sequence, known as the proto-spacer adjacent motif (PAM), is also predicted to be 5’-NGG’-3’ in *S. mutans*, as is the case in *S. pyogenes* [18]. Given the similarities, one possible approach to CRISPRi in *S. mutans* could be to repurpose the endogenous Cas9_Smu_ nuclease into a nuclease-dead version, while also using elements of the original interference system (*e.g.* sgRNA). A CRISPRi system built around Cas9_Smu_ could function more effectively in *S. mutans* than one based on Cas9_Spy_. Heterologous expression of *S. pyogenes* dCas9 can be toxic to certain bacteria [19,20], although problems of this nature were not observed in a recent CRISPRi study in *S. pneumoniae* [14]. Irrespective of *S. pyogenes* dCas9 toxicity, the dCas9_Spy_ could perform suboptimally compared to dCas9_Smu_ simply due to codon-bias. Lastly, developing an *S. mutans* CRISPRi system has the potential to add to the number of tools for genome editing that are currently available.

Motivated by a desire to functionally study essential genes in *S. mutans*, we created a CRISPRi tool assembled around dCas9_Smu_. We explored sgRNA specificity and PAM requirements, and compared interference activity with dCas9_Spy_. Having demonstrated good CRISPRi activity, we then constructed a library of sgRNAs against all putative essential and growth-supporting genes [10]. With this library we evaluated growth and certain phenotypes in a subset of strains with CRISPRi-silenced essential genes. Finally, we used the technique to systematically study the impact that depletion of L-rhamnose-containing cell wall polysaccharides has on cell division and pathogenicity in an insect model.

## Results

### Robust gene knockdown with a titratable dCas9_Smu_ system

More than 95% of the 477 *S. mutans* strains for which whole or draft genome sequences are presently available carry at least one CRISPR-Cas system (Fig 1A). The Type II-A CRISPR-Cas system, which can be repurposed for CRISPRi, is the most prevalent across the species with 339/477 strains carrying a complete system (Fig 1A). For reprogramming of *S. mutans* UA159 Cas9 (SMu.1405c) into a CRISPRi system, we began by mutating two amino acid residues (D10 and H840) known to be required for the nuclease activity of *S. pyogenes* Cas9 [21] (Fig 1B). We were able to confirm the DNA-binding and repressing activity of dCas9_Smu_ by expressing a sgRNA against a constitutively expressed *gfp* gene [22] (Fig. 1C). Next, the *S. mutans dcas9* gene was cloned into pZX9 [23], such that expression could be activated by xylose. Prior to this, we discovered, using green fluorescent protein (GFP) as a readout for xylose-induced gene expression, that the P_*xyl*_ promoter was less active when glucose was the sole carbohydrate source compared to maltose (Fig S1). We suspect the different expression levels on glucose versus maltose could be due to carbohydrate catabolite repression and/or effects of certain phosphoenolpyruvate:sugar phosphotransferase system (PTS) components on P_*xyl*_ activity. Thus, we used the defined media ‘FMC’ containing maltose for all CRISPRi related experiments. To make the CRISPRi strain (Fig 1D), the sgRNA plasmid was integrated in single copy into the *S. mutans* genome in a strain that contained i) a *gfp* gene driven by the constitutively active P23 promoter and integrated into the chromosome, ii) the P_*xyl*_-*dcas9*_Smu_ plasmid and iii) point mutations that prevent translation of SMu.1405c (*cas9*; start codon mutated to stop codon). *S. mutans* Cas9 must be mutated, otherwise endogenous *cas9* expression in combination with sgRNAs would likely cause lethal chromosome damage; *S. mutans* is unable to repair the double-strand cleavage catalyzed by a functional Cas9.

**Figure 1.**
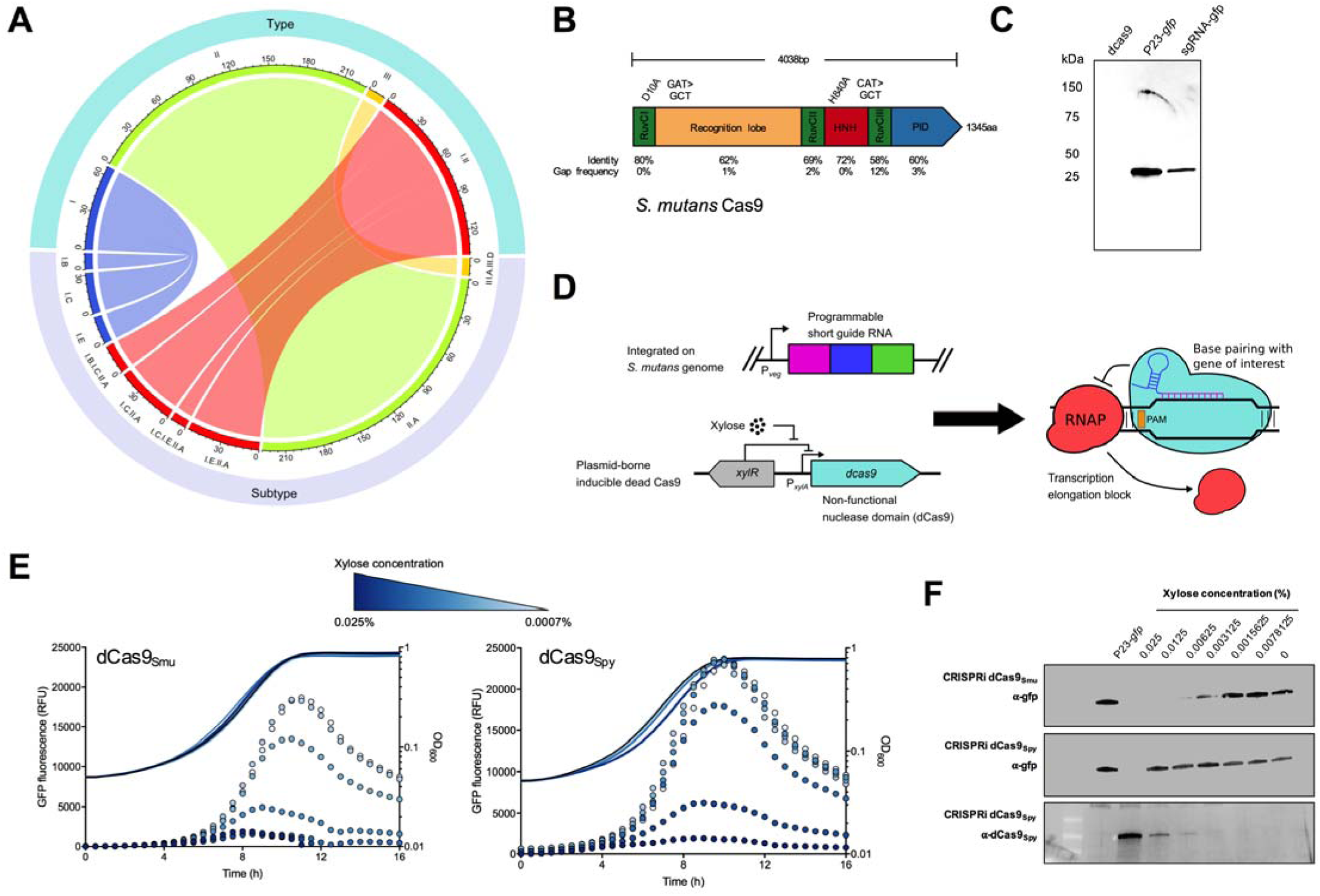
Design of an inducible CRISPRi gene-silencing system for *S. mutans*. (A) Distribution of CRISPR-Cas systems in *S. mutans* by type (top half) and subtype (bottom half). For example, 67 *S. mutans* strains contain Type I systems (blue), and the subtypes of this system are B, C, and E (I-C is the most common). The numbers on the exterior of the plot denote the number of *S. mutans* strains that possess a certain CRISPR-Cas system. (B) Diagram of the Cas9 protein of *S. mutans* UA159 showing domains. The two conserved amino acids required for nuclease activity in the *S. pyogenes* Cas9 protein that were mutated to create the *S. mutans* dCas9 protein are indicated. Each domain was compared with *S. pyogenes* Cas9 for percentage amino acid identity. (C) Gene-silencing activity of endogenous dCas9_Smu_ was assessed by expressing an sgRNA with complementarity to a chromosomally integrated *gfp* gene and comparing GFP protein levels using Western blotting (sgRNA-*gfp*). Also shown is a negative control, d*cas9* without *gfp*, and a positive control strain that constitutively expreses the *gfp* gene from the P23 promoter (see text for details). (D) Overview of the CRISPRi system engineered for *S. mutans*, which consists of two primary elements, a chromosomally located sgRNA and a plasmid-borne inducible dead Cas9. Combined, these two elements can block gene transcription as shown on the right. (E) Comparison of dCas9_Smu_ (left) and dCas9_Spy_ (right) *gfp* gene silencing activity using real-time fluorescence monitoring. Fluorescence readings are shown as dots, and cell density (OD_600_) readings as lines. As the xylose concentration was decreased, data points change from a dark to a light blue color. (F) Comparison of dCas9_Smu_ (top panel) and dCas9_Spy_ (middle panel) *gfp* gene silencing activity with Western blotting as as a function of xylose concentration. The bottom panel is a Western blot against the xylose-inducible dCas9_Spy_; as this antibody detected dCas9_Spy_, but not dCas9_Smu_. Western images are representative of three independent replicates.

Having developed the system, we moved to assessing its efficiency. Using the *S. mutans* CRISPRi technique, we observed titratable repression of GFP fluorescence (Fig 1E) and GFP protein levels (Fig 1F). Importantly, there were no obvious effects on the growth *S. mutans* as increasing amounts of xylose were added (up to 0.025% w/v). Also of note, slightly higher amounts of xylose (0.025% vs. 0.00625%) were required to achieve similar levels of repression of GFP production in a strain containing a xylose-inducible *S. pyogenes dcas9* (Figs 1E and F). We also observed slight growth impairments as increasing amounts of dCas9_Spy_ were produced (Fig 1E). Overall, the differences in performance of the two Cas9 constructs were minor. Next, we measured gene-silencing (absence of GFP fluorescence) at the single-cell level (Fig S2). The number of cells escaping CRISPRi-mediated gene knockdown (GFP positive) was very small, indicating that, at saturating amounts of xylose, the effects of the *S. mutans-*derived CRISPRi system are uniform across the population when the dCas9 protein is delivered on a multi-copy plasmid. The pZX9 plasmid that carries the xylose-inducible system uses the streptococcal replicon from pDL278. Without antibiotic pressure we have shown that pDL278 is stable in *S. mutans* for at least 50 generations [22].

RNA sequencing (RNA-seq) experiments were conducted to test the effects of the CRISPRi system on the *S. mutans* transcriptome. We compared the transcriptome of the CRISPRi strain without and with the addition of xylose (0.1%). With xylose, we observed an increase in dCas9 expression and a concomitant decrease in *gfp* expression compared to the no xylose condition (S1 Table). This was specific to the presence of sgRNA-*gfp* as we observed no reduction in *gfp* expression in a strain lacking the sgRNA and xylose added (no sgRNA in S1 Table). As has been noted by other authors [13,14] we measured a polar effect of CRISPRi on genes downstream of the target gene (*msmK* and *dexB* are downstream of the location of the *gfp* gene). Importantly, the effect of CRISPRi induction on the *S. mutans* transcriptome was minimal. We observed no *S. mutans* gene expression changes with a log_2_FC >2, and only twelve genes with a log_2_FC >1 (FDR <0.05). Furthermore, these twelve genes typically contained low read counts, indicating that the slight changes in gene expression could be attributable to noise. In the absence of CRISPRi, but with xylose (no sgRNA strain), no changes in *S. mutans* gene expression were observed; so 0.1% xylose has no discernable impact on the *S. mutans* transcriptome. From these RNA-seq results, we conclude that the CRISPRi system is highly specific and does not appreciably alter *S. mutans* gene expression other than targeted genes.

### Guide RNA design principles for the sgRNA-Cas9_Smu_ system

To begin to optimize the *S. mutans* CRISPRi system, we examined parameters that are important for functionality; the first of these was sgRNA specificity. sgRNA-Cas9 systems that operate with lower sequence stringency could be prone to off-target effects that occur when guide RNA and dCas9 bind to a site(s) in the genome that was not intended to be affected. To test this, we cloned an sgRNA with a complementary sequence for the gene *atpB*, which encodes the beta subunit of the F_1_F_O_ proton-translocating ATPase. We modified the sgRNA-*atpB* sequence by introducing substitutions along the length of guide RNA (Fig 2A). The sgRNA plasmids were added to competent *S. mutans* and plated onto selective agar. We expected that transformation into *S. mutans* of the sgRNA-*atpB* plasmid lacking sequence modifications would not be tolerated, leading to a very low transformation efficiency. As expected, transformation of the sgRNA-*atpB*^T^ (T; targeting the template strand) occurred at very low efficiency, 1 × 10^−5^ (Fig 2B). Conversely, transformation efficiency in a Δ*cas9* (deletion of Smu.1405c) strain, the dead *cas9* strain, or in a strain carrying a deletion of 20-bp of the *atpB*^T^ sequence from the sgRNA (sgRNAneg) was 100- to 1000-fold more efficient than with the intact *atpB* sgRNA (Fig 2B).

**Figure 2.**
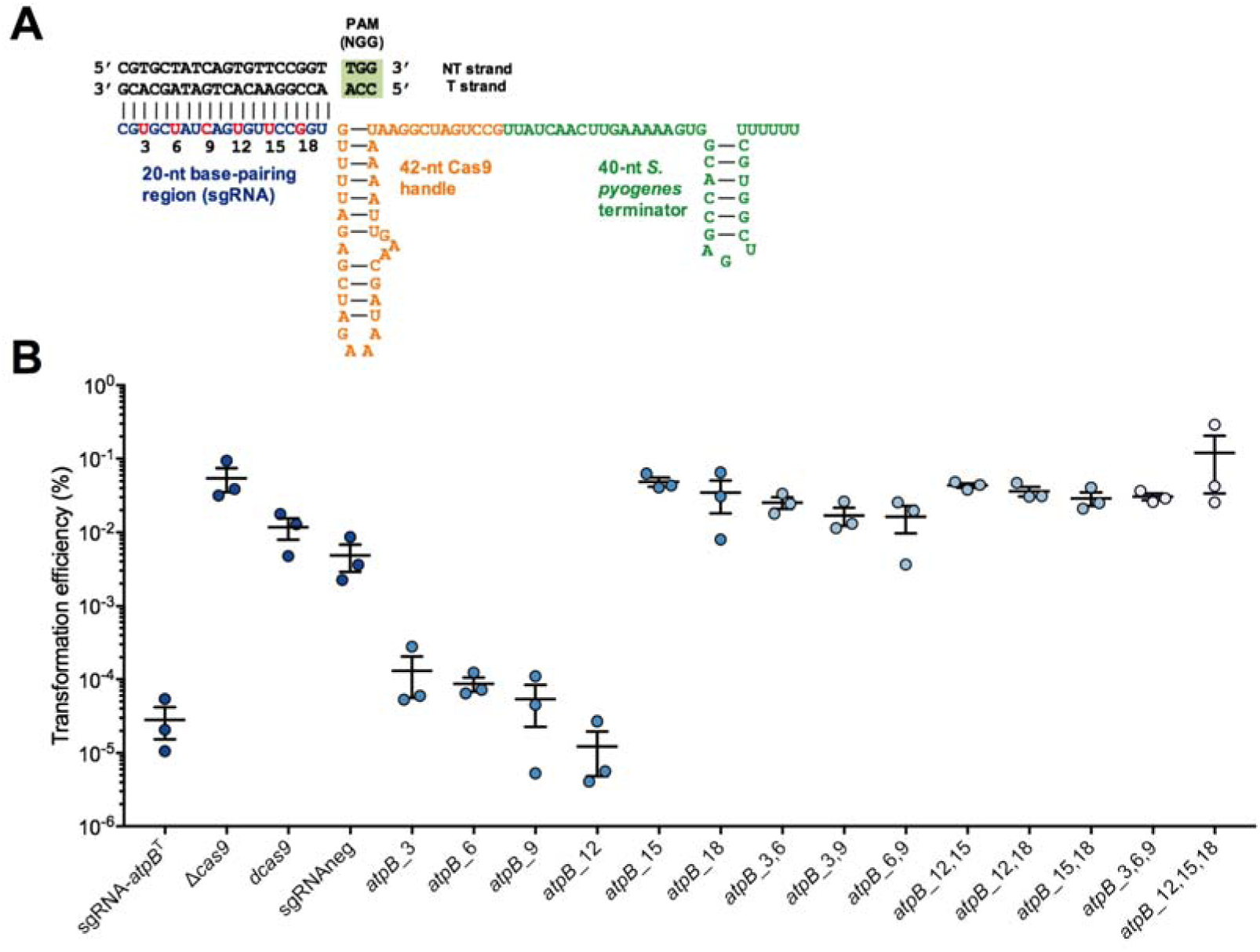
Determining guide RNA specificity of Cas9_**Smu**_. (A) Diagram of the sgRNA for *atpB* and the mutagenesis strategy. Positions 3, 6, 9, 12, 15 and 18 of the sgRNA were mutated (including in pairs or triplets) as a way of investigating sgRNA specificity (see text for additional details). (B) Plasmid interference assay of wild-type and mutated sgRNA-*atpB* plasmids, with Δ*cas9*, d*cas9* and sgRNAneg serving as controls (for no plasmid interference). If the plasmid interfered with *S. mutans atpB* expression, a lower transformation efficiency would be expected.

We next evaluated the effects of expressing sgRNAs with one, two or three base mutations within the *atpB*^T^ sequence. Single base-pair mismatches were tolerated at the 5’ end of the sgRNA (*atpB*_3, *atpB*_6, *atpB*_9 and *atpB*_12). Plasmids with mismatches closer to the PAM (*atpB*_15 and *atpB*_18) transformed with higher efficiency into *S. mutans*, indicative of a loss of sgRNA activity. All of the double and triple base mismatches reduced the activity of the sgRNAs, as these plasmids transformed *S. mutans* at levels comparable to the *cas9-*deficient strain (Fig 2B). These results suggest that near perfect sgRNA sequence complementarity is required to silence genes, and thus it should be possible to minimize off-target effects with careful design of sgRNAs.

Another important feature of CRISPR-Cas9 systems, both as a natural bacterial immune system and for CRISPRi, is the proto-spacer adjacent motif (PAM). This sequence is required for the Cas9 protein to recognize interference targets and is also critical during the acquisition of CRISPR spacers (for Type II-A systems) [21,24]. For a given bacterial strain, Type II-A PAM sequences are generated by aligning CRISPR spacers to phage genomes, extracting the sequence downstream of the spacer target, and then building a conserved sequence from these. The difficulty with taking this approach with *S. mutans* UA159 is that there are too few CRISPR spacers from which to build a reliable PAM sequence. Consequently, we compiled PAM sequences from a large (>450) collection of *S. mutans* genomes. After narrowing the data to strains that had Type II-A CRISPR-Cas systems and ≥5 spacers, PAM sequences were generated for 100 individual *S. mutans* strains using WebLogo (Fig S3). Per this bioinformatics analysis, a highly conserved PAM for Cas9_Smu_ was not evident. Rather, a significant number of *S. mutans* strains with Cas9 contained spacers that might require the 5’-NGG-3’ recognition sequence for phage DNA interference, and these spacer arrays grouped together by alignment (Fig 3A, yellow color). For other *S. mutans* strains with Cas9, PAM sequences were more random, but at least two other groups, 5’-TTN-3’ and 5’-NAAA-3’, existed (Fig S3). We observed no obvious grouping for *S. mutans* of the predicted PAM sequences by Cas9 protein identity (Fig 3A). We reasoned that PAM recognition might be protein sequence specific and thus expected that the different PAM groups predicted by bioinformatics would also group together. A lack of coherence between the predicted PAM sequences and Cas9 protein sequences suggests that the PAM sequences assembled via bioinformatics may not be the best indicator of the requirements for recognition for all *S. mutans* strains. So, to investigate DNA recognition in greater detail we developed an interference assay with GFP fluorescence as the readout. Using site-directed mutagenesis we incorporated assorted PAMs into a construct in which the P23 promoter could drive the expression of a *gfp* gene (Fig 3B). This assay confirmed that *S. mutans* dCas9 recognizes a 5’-NGG-3’ PAM (Fig 3C). With 5’-NAG-3’ we observed moderate CRISPRi activity with dCas9_Smu_, although the results varied as a function of the DNA base present in the first position, with up to an 80% loss of GFP fluorescence. Weak binding of *S. pyogenes* Cas9 to a 5’-NAG-3’ PAM has been reported before [25–27]. Nuclease activity via 5’-NGA-3’ recognition has previously been reported for Cas9_Spy_ [26,27] but we were unable to observe CRISPRi activity with either dCas9 protein with a 5’-AGA-3’ PAM. No interference was observed with any of the other PAM sequence combinations tested.

**Figure 3.**
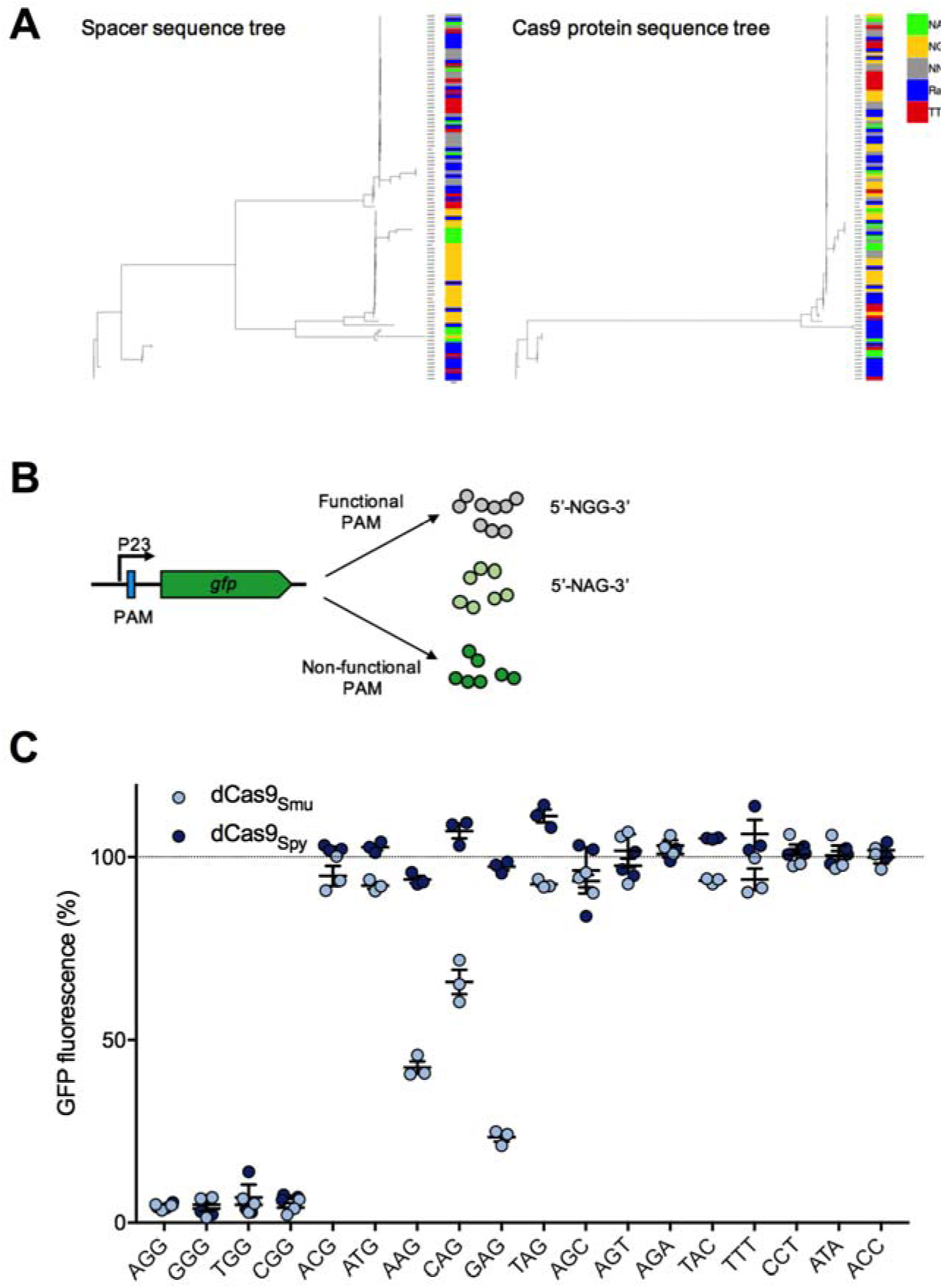
Investigating the proto-spacer adjacent motif (PAM) requirements of Cas9_**Smu**_. (A) Phylogenetic relatedness of extracted CRISPR-Cas9 spacer sequences (left) and Cas9 protein sequences (right). Consensus PAM sequences were built from spacer sequences and are available in Fig. S3. A key describing the colors chosen for consensus PAM sequences is shown on the far right. (B) To determine the optimal PAM sequence for dCas9_Smu_, different PAM sequences were inserted into between the P23 promoter and *gfp* gene. (C) The functionality of various PAM sequences were compared by measuring CRISPRi repression of GFP fluorescence for both *S. pyogenes* and *S. mutans* Cas9. Repression was calculated as a percentage of maximal fluorescence without exogenous xylose compared to fluorescence with 0.1% xylose added. Bars show the average and standard deviation, and light blue dots are the individual values, obtained using the dCas9_Smu_ construct, whereas the dark blue dots show data obtained using dCas9_Spy_. Maximal repression was measured with a 5’-NGG-3’ PAM, with moderate repression for some 5’-NAG-3’ PAMs.

### Creation of a comprehensive library of essential gene targeting sgRNAs

We constructed a library of sgRNAs targeting 253 potentially essential and growth supporting genes (Fig 4A). These genes (S2 Table) were selected from previously conducted Tn-seq experiments [10], Tn-seq performed as part of this study, and genes shown to be essential in previous studies (e.g. *vicR/walR*). We were unable to design sgRNAs for 21 putative essential or growth supporting genes, of which over half (12) were ribosomal proteins. These genes were small (average size of 275-bp) and generally lacked PAM sites that would give desirable sgRNA sequences (*e.g.* RNA folding, off-target effects, location in 5’ region of gene). During the first round of cloning (see Materials and Methods) we obtained correct sgRNA clones for 181 of 282 (64%) of the desired strains. After three rounds of cloning 259/282 (92%) of the sgRNAs were successfully cloned. We were unable to obtain sgRNA clones for 23 (8%) genes. After plasmid cloning in *E. coli* 10-beta, sequence-verified sgRNA plasmids were transformed into *S. mutans* Δ*cas9* P_*xyl*_-*dcas9*_Smu_. While it would be possible to clone the sgRNA plasmids directly into *S. mutans*, we wanted to minimize the number of passages in *S. mutans* as CRISPRi strains are known to acquire suppressors when guide RNAs are expressed in their hosts [14].

**Figure 4.**
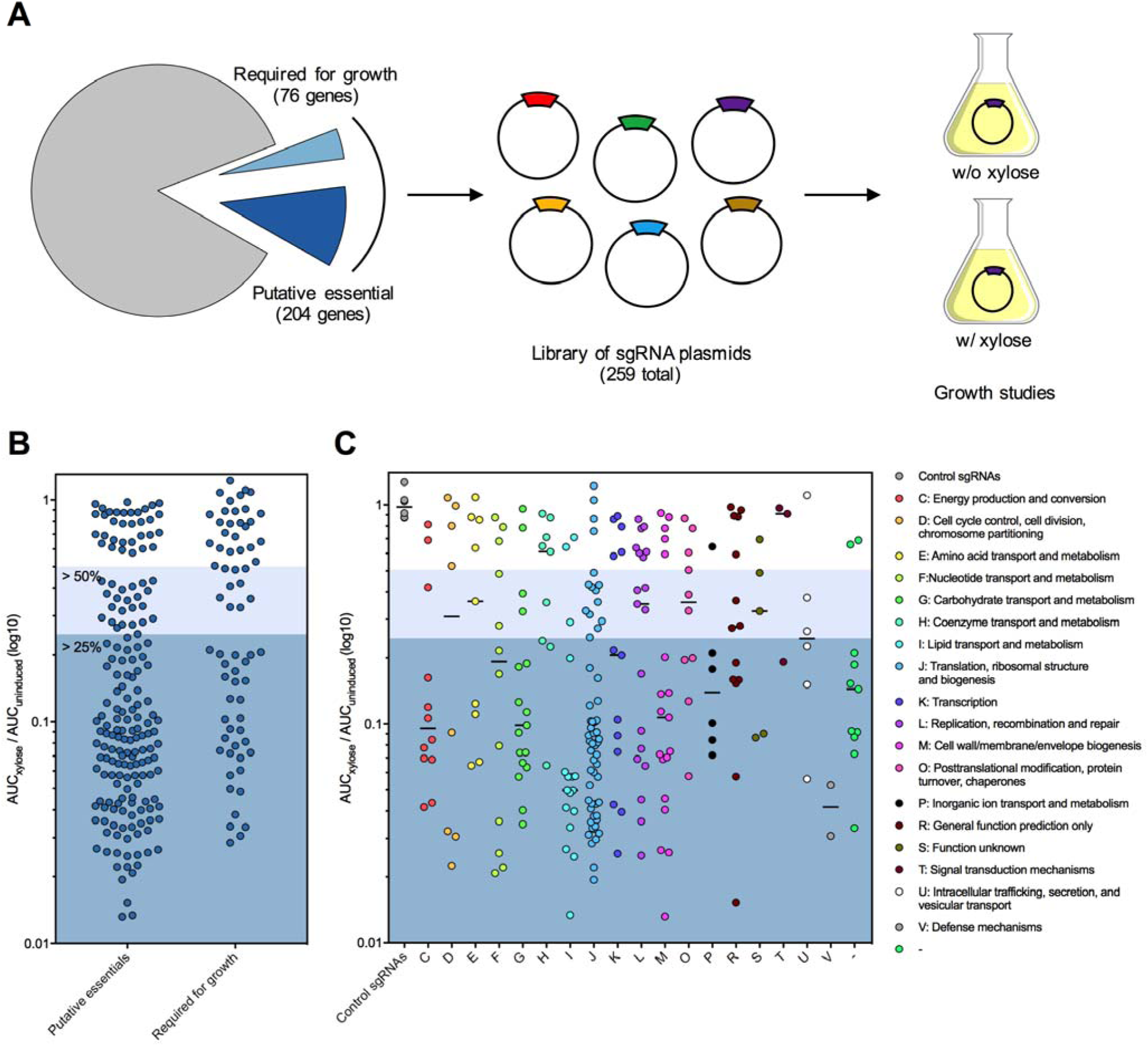
High-throughput growth analysis of a comprehensive library of sgRNAs targeting essential and growth supporting genes. (A) Overview of the CRISPRi growth study that included generating sgRNAs against >250 essential or growth-supporting genes. Growth analysis of CRISPRi strains (B) whether putative essential or required for growth and (C) sorted for physiological functions by Clusters of Orthologous Groups. CRISPRi strains were cultured in the presence or absence of xylose and the area under the curve (AUC) was calculated for each condition. Growth phenotypes were categorized according to the ratio of AUC_xylose_ to AUC_uninduced_ (the lower the number the stronger the growth defect).

### Growth phenotypes of strains with silenced essential genes

We measured the growth of each sgRNA strain in FMC-maltose either in the absence or presence of 0.1% xylose for 16 h using a Bioscreen C Automated Microbiology Growth Curve System. Afterwards, the area under the curve (AUC) was calculated for both conditions. Quantification of growth by AUC is useful as it combines growth phases (lag, exponential and stationary phases). For each strain, the AUC_xylose_ was divided by AUC_uninduced_. Growth phenotypes were grouped per these ratios: >0.5, minor phenotype; 0.25-0.49, moderate growth defect; <0.25, strong growth defect. It is important to note that the differences between strains slightly above or below the ratio cut-off points are not substantial. Growth defects were observed in 194/253 (77%) of the strains expressing sgRNAs that were designed to target essential or growth supporting genes (Fig 4B; S2 Table). Overall, we observed a higher percentage of growth defects for strains with silenced essential genes versus growth-supporting genes, 82% and 62%, respectively (Fig 4B). When comparing between different Clusters of Orthologous Groups (COGs), the two lowest median growth defects were observed for silenced genes in the translation (J) and lipid metabolism and transport pathways (I) (Fig 4C; silenced genes in the category defense mechanisms (V) displayed strong growth defects but we are unsure of these genes functions). The two highest median growth defects were observed when sgRNAs targeted genes related to coenzyme transport and metabolism (H) and signal transduction mechanisms (T) (Fig 4C). Growth defects were not observed for the six sgRNAs that targeted non-essential genes (*scrB, lacG, gtfB, bceA, gfp, gfpprom*; see S2 Table). There was no significant relationship between the growth defect we observed and the position of the sgRNA within the gene or the size of the target gene (Fig. S4).

### Assessing cell biology phenotypes of CRISPRi strains with microscopy

Next we evaluated the morphological phenotypes of eighty CRISPRi strains placed in the following functional categories: fatty acid biosynthesis/membrane biogenesis, peptidoglycan synthesis, glycolysis, DNA replication, transcription, translation, cell division, and ATP synthase activity. Using DAPI staining, we were able to visualize the bacterial chromosome, and with nile red we could detect the cell membrane. The cell morphology defects of essential gene-targeted strains were diverse (Fig 5). Defects included increased cell chaining, abnormally large cells, heterogeneously sized cells, cell lysis and anucleate cells (Figs S5-S12). The most commonly observed morphological phenotypes across all the pathways investigated were increased cell chaining (*e.g. pfk, accD, dnaE, rs3, atpC*) and heterogeneously sized cells (*e.g. pgi, glmM, dnaG, eftS, atpF*) (Fig. 5A). It is important to note that there were strains without obvious growth defects that displayed major cell morphology defects (*e.g. ftsX* and *glmS*). Repression of genes involved in the glycolysis pathway led to significant growth defects and larger/misshapen cells (Fig S5). When *pgm*, encoding phosphoglucomutase, was repressed, cells appeared to be somewhat rod-shaped, probably associated with altered or deficient biosynthesis of cell wall polysaccharides. When genes involved in peptidoglycan biosynthesis were repressed, the most common phenotype was the formation of singular large round cells (Fig S6). The percentage of cells displaying this phenotype varied, with it being most common in a strain with silenced *murM*, which in *S. pneumoniae* encodes a protein that incorporates amino acids into the side-chain of peptidoglycan [28]. Repression of genes involved in the conversion of acetyl-CoA to malonyl-CoA (*accACD* and *bccP*), an important step in fatty acid biosynthesis, led to excessive chaining of cells (Fig S7). Repression of fatty acid biosynthetic (*fab*) genes yielded minor phenotypic changes, whereas silencing of genes downstream of the FAS II pathway (*plsX* and *cdsA*) led to more obvious morphological defects. As anticipated, when genes involved in DNA replication were repressed, not all cells contained nucleoids (Fig S8). In addition to anucleate cells, certain CRISPRi strains with repressed chromosome replication also showed severe morphological defects (e.g. *dnaE, dnaG, priA*). CRISPRi repression of translation-related genes increased cell chaining (Fig S9). When the ribosomal proteins *rs3* (small ribosomal subunit) and *rl11* (large ribosomal subunit) were repressed, we observed anucleate cells that lacked a well-defined cell envelope. CRISPRi silencing of genes encoding subunits of the F_1_F_O_-ATPase led to strong growth defects, increased cell chaining and variability in nile red staining (Fig S10). Repression of transcription-related genes also led to increased cell chaining and attenuated growth (Fig S11). The cell shape and size of *rpoB-, rpoC-*, and *rpoZ*-silenced CRISPRi strains were severely impacted. When genes involved in cell division were repressed, strains displayed heterogeneity in cell size and shape (Fig S12). Notably, with repression of *divIC, ftsA, ftsL* and *ftsZ* we observed similar morphological changes. Cells appeared to be rod-shaped, except that they bulged at the cell septum, probably consistent with an inability of these cells to complete septation. FtsZ is a tubulin homologue that forms a Z-ring at the midcell that is important for scaffolding and cytokinesis [29]. FtsA, DivIC and FtsL also localize to the midcell (at different stages) during cell division [29]. Repression of *gbpB* (glucan-binding protein B), an ortholog of the *S. pneumoniae* peptidoglycan hydrolase gene, *pcsB*, led to cell division defects. Although specific peptidoglycan hydrolase activity has not been observed for GbpB, the cell division defects we observed, along with the observation that GbpB can be localized to areas of active peptidoglycan synthesis [30], provide support for the idea that it may act enzymatically to affect the structure of the cell wall. Taken together, microscopic analysis of CRISPRi strains from different pathways has revealed generalized phenotypes of essential gene depletion, but also clear functional links between genes in established pathways.

**Figure 5.**
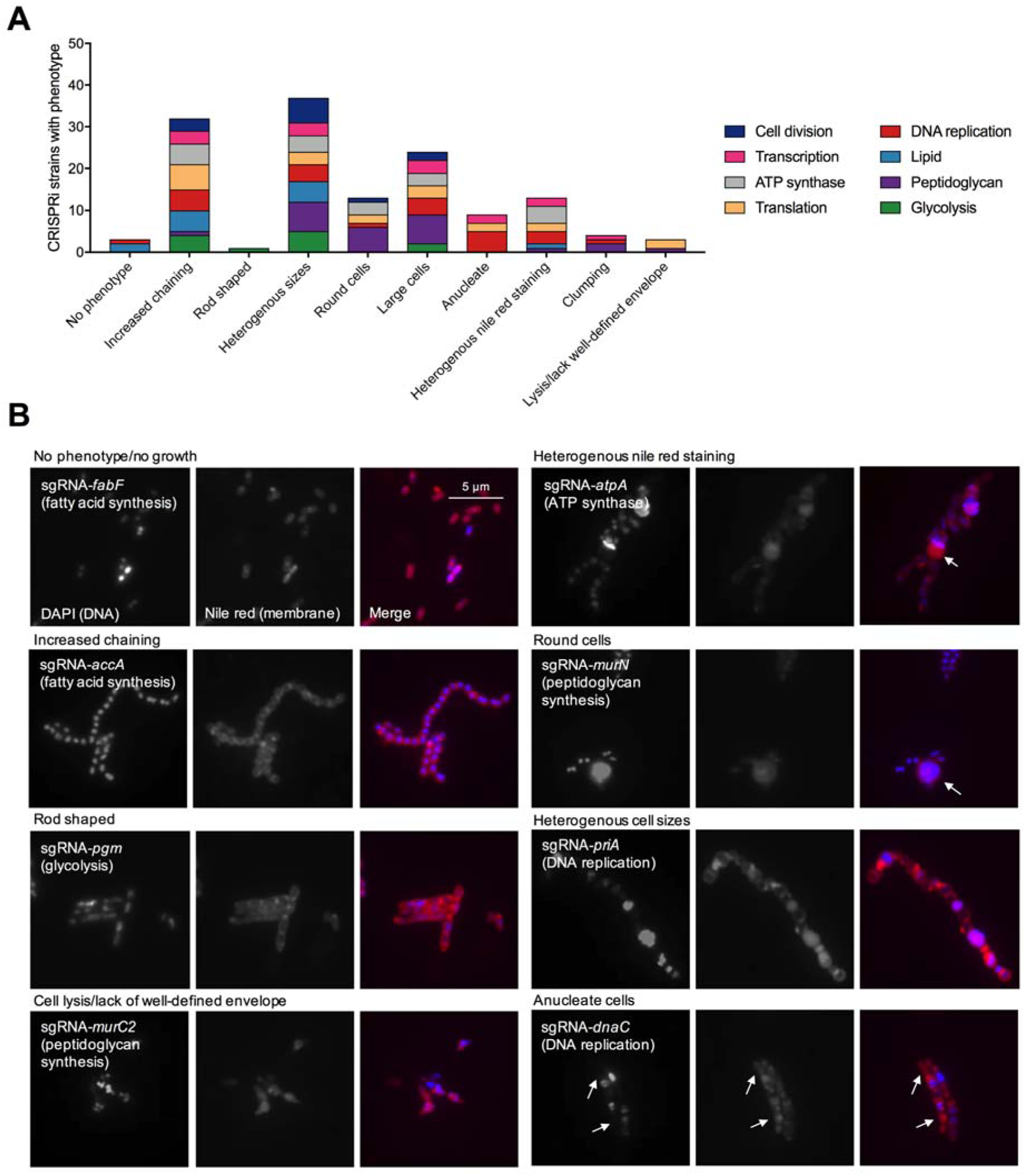
Microscopic analysis of the morphological defects caused by CRISPRi essential gene depletion. (A) CRISPRi strains, by functional pathway, were grouped according to cell morphology phenotypes. Strains with depleted essential genes can have more than one morphological defect (*e.g.* increased cell chaining and heterogeneous sizes). (B) Representative micrographs for several of the cell morphology phenotypes that were observed are shown. Additional microscopy across all the pathways investigated can be found in Figs. S5-S12. For sgRNA-*atpA* the arrow shows an area of increased nile red staining compared to other cells. The large round cells observed when silencing peptidoglycan biosynthesis is depicted with the sgRNA-*murN* micrograph. Arrows indicate potential anucleate cells when targeting the DNA replication-related gene *dnaC*.

### Identification of GpsB (SMu.471) as a critical cell division determinant of *S. mutans*

Next we wanted to employ CRISPRi as a tool to study poorly annotated essential genes in *S. mutans*. The gene SMu.471 and the intergenic region downstream of it were of interest to us as we noticed that this region did not tolerate transposon insertions (Fig 6A). After conducting BLASTn analysis of the intergenic region, we realized that the RNA component of RNase P, *rnpB*, is located here. The *rnpB* transcript, as measured by RNA-seq, was highly abundant, which is consistent with what has been observed in other bacteria [31]. RNase P is a ribonucleoprotein that catalyzes the maturation of the 5’ end of pre-tRNAs [32]. To characterize the gene SMu.471, we began with CRISPRi phenotyping, which showed that depletion leads to a strong growth defect (Fig 6B). Depletion of SMu.471 also led to cell morphology defects (Fig 6D). Cells were wider and larger than wild-type and some appeared to not be correctly dividing (Fig 6D). Although annotated as a hypothetical protein, BLASTp analysis showed that SMu.471 contains a GpsB/DivIVA domain. SMu.471 aligned best with GpsB proteins from other Gram-positive organisms (Fig S14), consistent with it being smaller (112 aa) than true DivIVA proteins. Of note, the *S. mutans* UA159 genome contains three proteins with DivIVA domains, SMu.471, SMu.557, and SMu.1260c (Fig S14). SMu.557 (271 aa) is larger than SMu.471, whereas SMu.1260c is the smallest protein of the three (59 aa) (Fig S14). GpsB proteins are smaller than DivIVA proteins because of a reduction in the length of the C-terminal domain (CTD). GpsB has been shown to interact with a diverse assortment of proteins involved in cell division, with penicillin-binding protein (PBP) interactions (predominately class A) being most common [33]. We tested the interaction of GpsB with PBPs, as well as with the cell elongation protein MreC [34–36], or the serine/threonine protein kinase PknB [34], using a bacterial-two-hybrid assay. We observed self-interaction of GpsB and a weak interaction with PBP1a, but no interaction was evident for either MreC or PknB (Fig 6E). Collectively, the CRISPRi growth defect, cell morphology changes and the interaction with PBP1a suggest that GpsB is a critical cell division protein in *S. mutans*.

**Figure 6.**
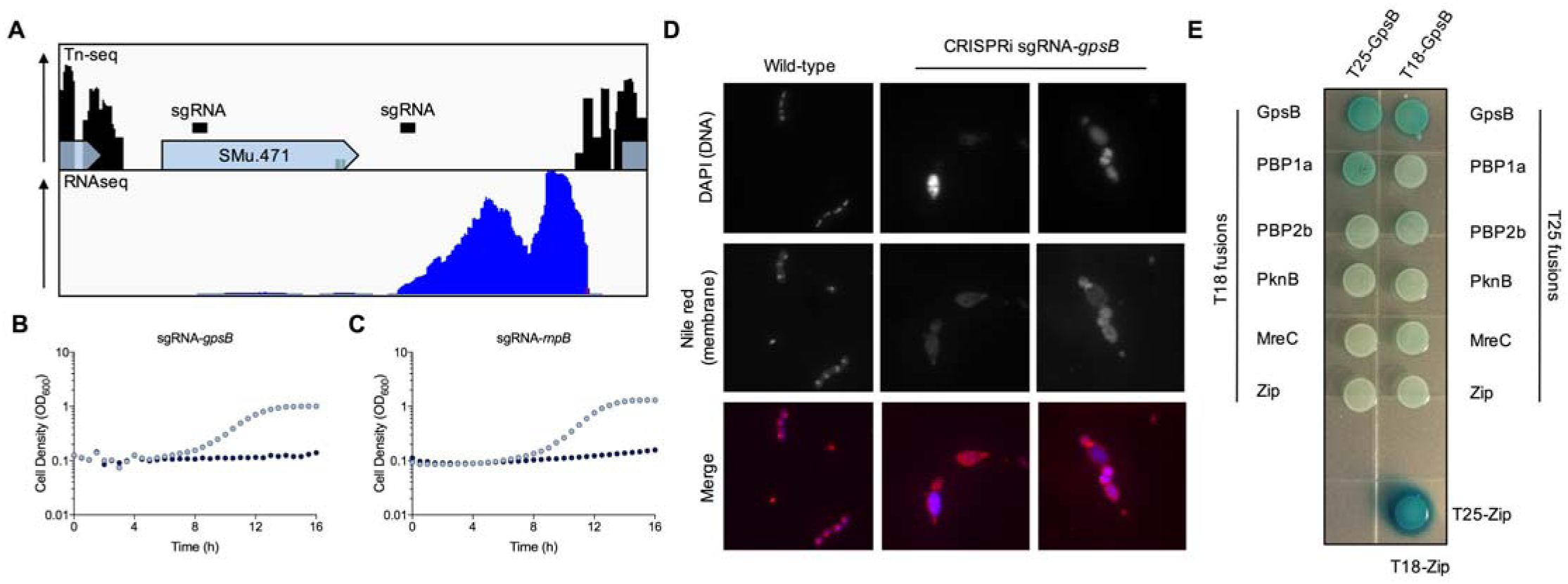
Identification of the essential peptidoglycan synthesis regulator GpsB in *S. mutans*. (A) Distribution of Tn-seq reads (top panel) within SMu.471 and the downstream intergenic region. Each peak corresponds to a transposon insertion with a lack of peaks within a gene representative of an essential gene (*e.g. SMu.471)*. RNA sequencing reads (bottom panel) obtained from mid-log *S. mutans* cells grown in FMC-maltose. High levels of transcription are evident in the region downstream of SMu.471, which is likely to be the RNA component of RNase P (*rnpB*). CRISPRi growth phenotypes of silenced *gpsB* (SMu.471; B) and *rnpB* (C). Strains were cultured in FMC-maltose without (light blue dots) or with (dark blue dots) 0.1% xylose. (D) Microscopic analysis of a CRISPRi strain with depleted levels of *gpsB*. Strains were cultured overnight in FMC-maltose containing 0.1% xylose, stained with DNA and membrane staining dyes, and then imaged with fluorescence microscopy. Images are representative of five independent replicates. (E) Bacterial two-hybrid screening to test interactions between GpsB and putative interaction partners. Interactions were tested using constructs cloned into two plasmids pKT25 and pUT18C (T18 and T25 fusions). A positive control is shown on the bottom right (T25-zip with T18-zip). GpsB interacts with itself and has a weak interaction with PBP1a as shown by blue colonies (β-galactosidase activity has been restored by interactions between the two proteins; see S1 file for a description of the methodology).

### L-rhamnose glycopolymers are critical for *S. mutans* cell division and pathogenicity

Having shown that CRISPRi allows for the functional study of essential genes, we next focused our attention on a pathway that is thought to be of major importance to *S. mutans*, rhamnose-glucose polysaccharide (RGP) biosynthesis. Certain streptococcal species lack cell wall teichoic acids (WTAs) and instead produce RGPs that are covalently linked to the peptidoglycan [37]. RGP biosynthesis begins with a four-step pathway encoded by the genes *rmlABCD* that converts glucose-1-phosphate into dTDP-L-rhamnose. The last gene in the pathway, *rmlD*, resides in an operon (SMu.824-835) that contains genes required for additional RGP biosynthesis steps and transport of the carbohydrates through the cell membrane. By our Tn-seq analysis, most of the genes involved in producing RGP are either essential or required for the growth of *S. mutans* UA159 (Fig 7A) [10]. L-rhamnose biosynthetic pathways are also essential in *S. pyogenes* [38] and *Mycobacterium smegmatis* [39]. Several of these genes have, however, been mutated in *S. mutans*, including *rmlBC* [40], *rmlD* [38], *rgpE* [30,41], *rgpF* [41,42], and *rgpG* [43]. Inconsistencies between our observations and others could be related to the nature of our Tn-seq selection and/or the presence of undetected suppressor mutations that bypass essentiality in strains that carry RGP gene deletions in other studies. Use of CRISPRi to study RGP function facilitates the rapid evaluation of depleting each gene product under conditions that allow for selective pressure to be applied only when xylose is provided in the medium. By CRISPRi, down-regulation of every gene directly involved in RGP biosynthesis displayed severe growth defects when silenced with sgRNAs (Fig 7B; Fig S13). In the invertebrate infection model organism *Galleria mellonella*, strains with silenced *rmlB, rgpC*, and *rgpF* genes displayed attenuated virulence compared to wild-type *S. mutans* UA159 and a d*cas9* strain (Fig 7C). For this proof-of-concept study, sgRNAs were expressed in an *S. mutans* strain where the endogenous Cas9 had been converted to dCas9. Using this approach, no xylose is required and instead genes are repressed by the constitutively produced dCas9 (Fig 1C). A strain with silenced *rmlD* was not attenuated. However, we suspect this strain acquired suppressor mutation(s) as it originally grew very poorly, but after some passage it was able to grow to a final OD_600_ similar to the parental strain. Having shown that RGP biosynthesis genes are required for growth in laboratory media and during infection, we next investigated their importance to *S. mutans* cell biology. Consistent with previous observations [30,43], cellular morphologies were exceptionally aberrant, compared to the wild type (Fig S13). Cells were large, not consistently ovococcoid, and formed longer chains than normal. Although resolutions were limiting, it also appeared that certain large cells contained multiple nucleoids by DAPI strain (Fig S15). To gain greater ultrastructural insights, we investigated cell biology phenotypes of *rmlB, rmlD, rgpC*, and *rgpF* silenced strains with transmission electron microscopy (TEM). In these strains, division site positioning was irregular and not always in the mid-point of cells (Fig 7D). There were instances where the division septum was unable to traverse the cell (*rmlB, rmlD, rgpF*) or cells had double septa (*rgpF*) (Fig 7D). Accordingly, large cells with incomplete septation appeared to have multiple unequally sized compartments, which likely has a deleterious effect on chromosome segregation, and might explain the unusual patterns of DAPI staining observed during fluorescence microscopy (Fig S15). In *rgpC*-silenced cells, the equatorial ring, an outgrowth of cell wall material that is normally duplicated and separates as cells divide, was abnormally thick and in some cases was ‘peeling’ back from the cell (Fig 7D). Measurements from the CRISPRi strain micrographs using ImageJ confirmed that they were wider than wild-type cells (Fig 7E).

**Figure 7.**
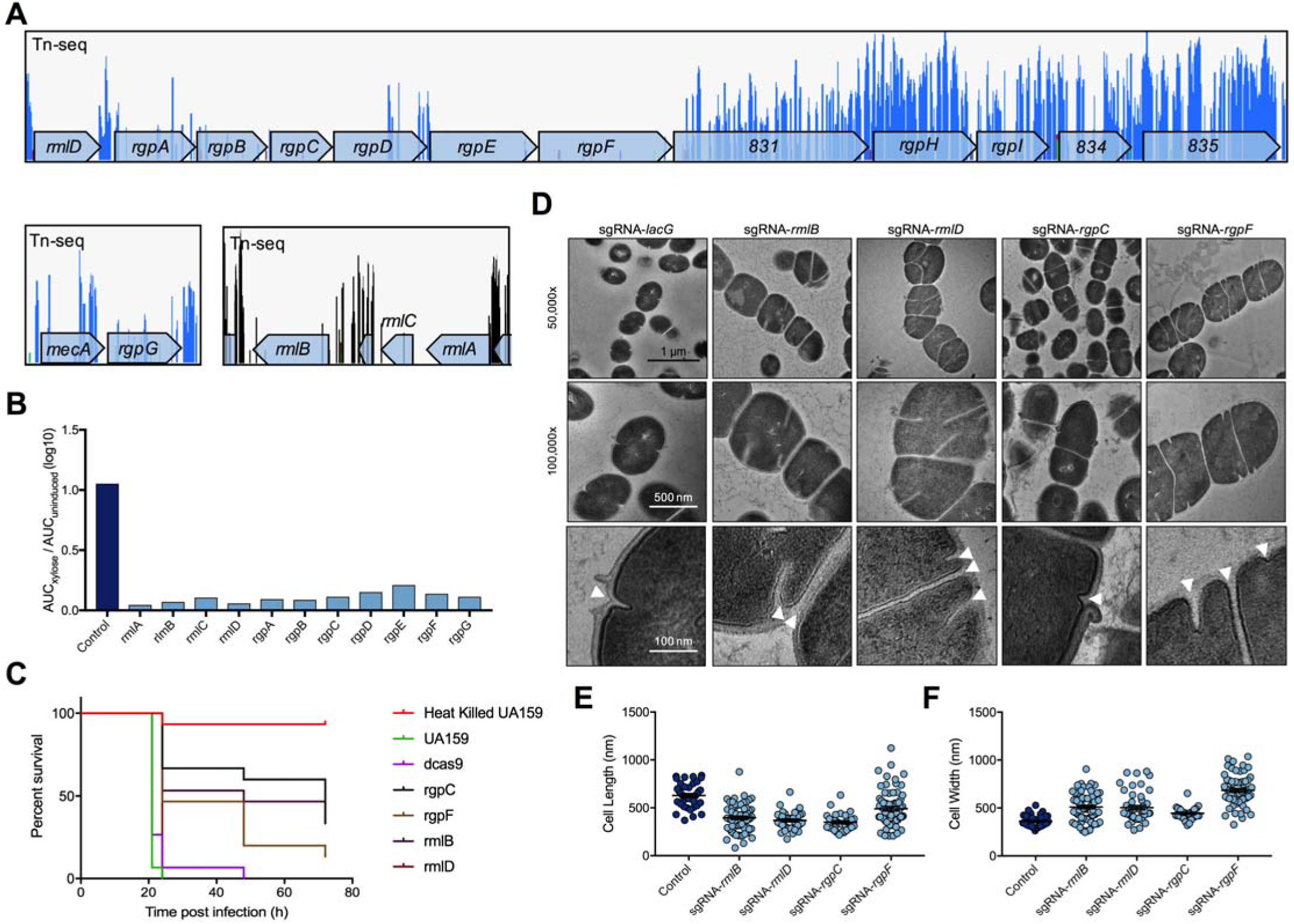
Rhamnose-glucose polysaccharide biosynthesis genes are required for growth, **pathogenesis and correct cell division of *S. mutans***. (A) Distribution of Tn-seq reads within genes that function in RGP biosynthesis (*rmlA*-*rmlC, rgpG*, and *rmlD*-*rgpF*; SMu.831-835 reside in the operon that starts with *rmlD*). Each peak corresponds to a transposon insertion with a lack of peaks within a gene representative of an essential gene (*e.g. rmlA* or *rgpE*). (B) Growth phenotypes of silenced RGP biosynthesis genes compared to a control strain (sgRNA-*lacG*). (C) Pathogenesis phenotypes of silenced RGP biosynthesis genes (*rmlB, rmlD, rgpC*, and *rgpF*) in a *Galleria mellonella* infection model. Heat killed UA159 served as a negative control, and wild-type UA159 and d*cas9* absent sgRNA as positive controls. (D) Electron micrographs of CRISPRi strains targeting the same genes investigated in the pathogenesis model. Arrowheads point to putative division septa. Fluorescence microscopy images and growth curves for CRISPRi strains for all of the genes involved in this pathway can be found in Fig. S13. Cell length (E) and width (F) measurements from TEM micrographs using ImageJ.

## Discussion

Essential genes represent promising targets for antimicrobials. The essential genome of *S. mutans* is comprised of ∼300 genes and, like other bacteria, is enriched for genes that function in processing genetic information, energy production, and maintenance of the cell envelope. However, until the recent development of CRISPRi technologies, high-throughput functional studies of these genes has not been possible. Here we demonstrate the conversion of the *S. mutans* Cas9 nuclease into a gene silencing tool for the systematic interrogation of the essential genome. Our system is relatively streamlined as it only requires a constitutively expressed sgRNA and a xylose-inducible dCas9. Importantly, it allowed us to study all the known essential genes in a relatively short time. Most of the genes predicted to be essential by Tn-seq displayed strong growth defects once silenced by CRISPRi. Furthermore, CRISPRi strains with sgRNAs targeting essential genes displayed morphological defects when analyzed by microscopy. This work confirms the importance of these genes to the integrity of *S. mutans* cells, and of their potential as interesting therapeutic targets for the control of *S. mutans* or certain other pathogenic streptococci.

One interesting finding of this work is that Cas9_Smu_ recognition of target DNA shares similarities with Cas9_Spy_. First, guide RNA specificity of Cas9_Smu_ is similar to Cas9_Spy_ as single base-pair mismatches are not tolerated in the 5’ region of sgRNA (the seed region) [44,45]. Cas9_Smu_ was also unable to cause DNA cleavage *in vivo* when double or triple base-pair mutations were introduced into the sgRNA sequence, again consistent with findings in Cas9_Spy_ [45]. Second, PAM recognition relied on a 5’-NGG-3’ sequence. We also observed moderate CRISPRi activity with a 5’-NAG-3’ PAM. Cas9_Spy_ activity also relies on a 5’-NGG-3’ PAM with weak DNA recognition reported for the non-canonical PAMs 5’-NAG-3’ and 5’-NGA-3’ [25–27], although we were unable to measure this with our assay. Although target recognition also relies on other factors [46], we are confident that off-target effects should be minimized when using a 5’-NGG-3’ PAM and a guide RNA designed to ensure a lack of complementarity (two or more mismatches) to other regions of the genome. The similarities of Cas9_Smu_ and Cas9_Spy_ are interesting given that other streptococcal Cas9 nucleases require PAMs that differ to 5’-NGG-3’ [20,47]. For example, the Cas9 of *Streptococcus canis*, which shares ∼90% sequence similarity to Cas9_Spy_, recognizes a 5’-NNG-3’ PAM [47]. Bioinformatics analysis of the species-wide PAM requirements were cryptic, but it is possible that other *S. mutans* Cas9 proteins recognize alternative PAMs *e.g.* 5’-NAAA-3’. Intriguingly the Cas9 of *Streptococcus macacae* NCTC 11558 is presumed to recognize an adenine enriched PAM [48]. There is an arms race between the bacteria that generate Cas9 proteins recognizing new PAMs and the phage that undergo mutations to avoid CRISPR interference, with the latter being a confounding issue with PAMs that are generated solely using bioinformatic tools. Despite these limitations, it is possible there is species-wide PAM diversity, similar to what has been observed with *Neisseria meningitidis* [49]. This is an interesting area of study as it should aid in the engineering of new variants of Cas9 that recognize different PAMs, thus expanding the genetic engineering capabilities of the Cas9 proteins.

The CRISPRi system described here was able to perturb the expression of over 250 putative essential and growth-supporting genes. The severe growth defects exhibited by most of these strains emphasizes their critical functions in *S. mutans*. The percentage of silenced genes that displayed significant growth defects was comparable with what was observed for *S. pneumoniae* [14]. As noted above, CRISPRi has many advantages over more traditional approaches for studying essential genes, but the technique has its limitations. For example, failure to display an aberrant growth phenotype could be caused by sub-optimal sgRNA design or selection for suppressor mutations that bypass essentiality. Likewise, certain essential genes could still function with a low number of protein molecules per cell if the target gene is not completely silenced. In some cases, strains that displayed relatively minor growth defects when the target genes were depleted had significant alterations in cell morphology. Tn-seq, which was used to identify to candidate genes for this study [10], is a competitive assay, so even mutants with minor growth phenotypes could be significantly underrepresented at the chosen end-point. In this study, we supplemented our previous Tn-seq essential gene analysis [10] with a different Tn-seq approach, which involved plating the original library on BHI instead of blood agar, then passaging the libraries in BHI or FMC for ∼10 generations, as compared with ∼30 generations in our previous study [10]. The differences between the older and the newer Tn-seq analysis were minor (S2 Table). Overall, the CRISPRi growth study paired with comprehensive Tn-seq enquiries provide a solid foundation for determining which genes are essential in *S. mutans* UA159. It is reasonable to predict that some of these gene products could be targeted for the therapeutic control of *S. mutans* and possibly some other pathogenic streptococci.

In addition to growth studies, we were able to observe morphological phenotypes of CRISPRi strains with light and electron microscopy. Several morphology phenotypes were evident that were consistent with other high-throughput microscopic studies of essential gene depletions [14,15]. The changes in morphology could represent more generalized responses to an inability to grow or, in some cases, may prove useful in linking gene products that function in concert. For example, *ftsA, ftsZ, ftsL* and *divIC* shared similar phenotypes, consistent with their products being required for correct cell septum formation. Perturbation of peptidoglycan synthesis led in some cases to the formation of large round cells. Blocking or deleting peptidoglycan synthesis genes in other bacteria can lead to the generation of giant cells [50] or L-forms [51], although the phenotypes in this study appear distinct from those previously described variants.

The rhamnose-glucose polysaccharide (RGP) of mutans streptococci were used in the past to differentiate serotypes of strains that were previously all designated as *S. mutans.* Today, it is recognized that the species *S. mutans* consists primarily of Bratthall serotypes *c, e*, and *f*, along with the unusual serotype *k* strains [52,53]. In contrast, what was formerly designated serotype *b S. mutans* is now classified as *Streptococcus ratti*. It is also now well recognized that RGP is an important constituent of the envelope of mutans streptococci. Clear impacts on growth, pathogenicity and cell division were observed in strains that had silenced genes contributing to RGP biogenesis. The cell morphology phenotypes were stark, with depletion strains clearly exhibiting aberrant septum formation, as has been noted before in *S. mutans* and other related species [30,54,55]. An inability to divide normally has been linked to aberrant localization of the peptidoglycan hydrolase PcsB (GbpB in *S. mutans*) [30,55]. When *gbpB* was targeted with CRISPRi, cells appeared to be unable to divide correctly (Fig S12), but the phenotype did not match that associated with *rgp* gene silencing. Given the vital role and essentiality of the RGP biosynthetic pathway, coupled with the decreased virulence of certain *rgp* knockdowns reported here, RGP biogenesis may be an attractive antimicrobial target. Further, if RGP and peptidoglycan synthesis are linked, it is conceivable that targeting RGP biosynthesis could potentiate the efficacy of β-lactam and other cell wall-active antibiotics. Other attractive aspects of the RGP pathway as an antimicrobial target include the species or strain-specificity of the glycopolymers and the absence of the pathway in humans [37].

In summary, this is the first study to utilize CRISPRi in *S. mutans* as a genetic tool. We did so by repurposing the Cas9 that is widely distributed among the species, opening new avenues for investigating the diversity of CRISPR-Cas in *S. mutans* and other oral streptococci, and the possible development of next-generation CRISPR-Cas tools. We demonstrate that the technique provides a powerful approach for studying essential genes and gene function, in agreement with studies using other organisms [14,15]. We anticipate that these tools can be used to investigate new aspects of *S. mutans* cell biology and to identify promising therapeutic candidates.

## Materials and methods

### Bacterial strains and growth conditions

Strains of *Streptococcus mutans* were cultured in Brain Heart Infusion (BHI) broth (Difco) or chemically defined medium (FMC) [56] supplemented with 0.5% maltose. Streptococci were grown in a 5% CO_2_ aerobic environment at 37°C, unless stated otherwise. Strains of *Escherichia coli* were routinely grown in lysogeny broth (LB) with slight modifications (Lennox LB; 10 g/L tryptone, 5 g/L yeast extract and 5 g/L NaCl) at 37°C with aeration. Antibiotics were added to growth media at the following concentrations: kanamycin (1.0 mg/ml for *S. mutans*, 50 µg/ml for *E. coli*), erythromycin (10 µg/ml for *S. mutans*, 300 µg/ml for *E. coli*), spectinomycin (1.0 mg/ml for *S. mutans*, 50 µg/ml for *E. coli*) and ampicillin (100 µg/ml for *E. coli*). Strains and plasmids are listed in S3 Table.

### CRISPR-Cas bioinformatics

The 477 *S. mutans* genomes available in the National Center for Biotechnology Information (NCBI) GenBank in June 2018 were mined using CRISPRone [57] to determine the occurrence and diversity of CRISPR-Cas systems in this species. CRISPR arrays were also identified using CRISPRone and these were used to predict the PAM sequence based on spacer-protospacer matches using CRISPRTarget [58]. Afterwards, consensus PAM sequences for each strain were built using the WebLogo server [59].

### Construction of the xylose-inducible CRISPRi system

Xylose-inducible plasmids were cloned by following a protocol developed by Xie *et al* [23]. Three plasmids were generated that contained *gfp, dcas9*_Smu_ and *dcas9*_Spy_ inserts cloned into the pZX9 plasmid. For each plasmid, two linear fragments were created by PCR using pZX9 or *S. mutans* UA159 genomic DNA as templates. *dcas9*_Spy_ was amplified from a plasmid obtained from Addgene (Addgene #44249). *dcas9*_Smu_ was amplified from the strain described in the supplemental material. The *gfp* gene was amplified from previously constructed *gfp*-reporter strains [22]. The PCR fragments are designed so that they have overlapping complementary regions that facilitate concatemer formation during a 2^nd^ PCR. The products generated by this 2^nd^ PCR are transformed directly into *S. mutans*. Primers pZX9seqF and pZX9seqR were used to amplify gene inserts followed by Sanger sequencing. For cloning of the sgRNA plasmid (pPM∷sgRNA), a synthetic DNA fragment was generated (Integrated DNA Technologies) that contained the following components sequentially: 1) a P_*veg*_ promoter [22], 2) a 20-nt sgRNA targeting *gfp*, 3) a 42-nt dCas9 binding handle sequence, and 4) an *S. pyogenes* terminator. Restriction enzyme recognition sequences for *Sac*I and *Sph*I were included at the 5’ and 3’ ends of the sgRNA construct respectively. The synthetic DNA fragment was subsequently cloned into the pPMZ plasmid which integrates at the *phnA*-*mtlA* locus of *S. mutans* UA159. This plasmid was also used to generate plasmids containing sgRNAs that target other areas of the *S. mutans* genome. Protocols for cloning of additional strains are provided in S1 File. Oligonucleotides used in this study are listed in S4 Table.

### Green fluorescent protein CRISPRi assay

To test the CRISPRi system and compare the activity of dCas9_Smu_ to dCas9_Spy_ we used a GFP based assay. CRISPRi strains were cultured overnight, grown to an OD_600_ = 0.5, and then diluted 1:100 into FMC-maltose with or without xylose. For these assays the concentration of xylose was serially diluted. In triplicates, 200 µL samples were added to clear-bottomed, black-sided, 96-well microtiter plates. After loading of the cultures, the 96-well plate was placed in a Synergy HT microtiter plate reader (BioTek). The fluorescence and optical density (absorbance at 600□nm) were measured at 30-min intervals using Gen5 software (BioTek). The fluorescence settings were excitation at 485□nm, emission at 525□nm, and a sensitivity of 65. Data readings were collected, and the background fluorescence or OD600 was subtracted prior to data visualization using GraphPad Prism (v7) software (GraphPad Software). A slightly modified version of this assay was used for PAM comparisons (S1 File). The protocol for Western blotting of GFP and dCas9_Spy_ is available in S1 File.

### sgRNA library design and cloning

sgRNA sequences (20-nt) were selected with CRISPy-web [60]. With this tool we uploaded the *S. mutans* UA159 GenBank file (NC_004350.2) and selected essential genes. For each gene, the tool designs sgRNAs with predicted off-target matches (13-bp upstream of the PAM; off-targets containing 0, 1, 2, or 3-bp mismatches). To minimize off-target effects we only selected sgRNAs with two or more mismatches, which we have shown to diminish sgRNA activity (Fig 2B). We also aimed to select sgRNA sequences as close to the start of each gene as possible. For each sgRNA we predicted the secondary structure (20-nt sequence and 42-nt dCas9 binding handle) with the RNAfold program provided by ViennaRNA [61]. Only sgRNAs that maintained the hairpin structure of the dCas9 binding handle were selected for cloning. sgRNA sequences were cloned using the Q5^®^ Site-Directed Mutagenesis Kit (New England Biolabs). Primers were designed using the NEBaseChanger™ tool (v.1.2.9), substituting the sgRNA-*gfp* 20-nt sequence for whichever sgRNA we were cloning. Using the designed primers, the pPM∷sgRNA plasmid was amplified and then added to a mix of Kinase-Ligase-*Dpn*I for 5 min at room temperature. This mixture (5 µL) was then transformed into chemically competent *E. coli* 10-beta with selection on LB-kanamycin. For each sgRNA plasmid transformation, two independent clones were selected and cultured overnight. With each culture we made a glycerol stock for storage at −80°C, and purified plasmid with the QIAprep Spin Miniprep Kit (Qiagen). Each plasmid was Sanger sequenced using the pRCS1seqR primer (S4 Table).

### sgRNA plasmid interference assays

sgRNA sequence specificity was tested using a plasmid interference assay. A 20-nt sequence complementary to the template strand of the *atpB* gene (SMu.1528) was introduced into the pPM∷sgRNA plasmid with the Q5^®^ Site-Directed Mutagenesis Kit. Using the same kit, single, double, or triple base-pair mismatches were introduced into the 20-nt *atpB* sgRNA sequence. These plasmids were transformed into competent *S. mutans* and cells were serially diluted onto BHI agar, or BHI agar containing kanamycin. The percentage of cells that were transformed was calculated from the colony forming units present on both types of agar.

### CRISPRi growth assays

Strains of *S. mutans* carrying sgRNA cassettes were grown overnight in BHI with the appropriate antibiotics in a microaerophilic environment at 37°C. The following day the bacterial cultures were washed once in FMC-maltose, and then diluted 1:1000 into FMC-maltose or FMC-maltose containing 0.025% xylose. Triplicates of 300 µL were added to a Bioscreen C 10 × 10 well plate for each condition. OD_600_ was measured every 30 min for 16 h with a Bioscreen C Automated Microbiology Growth Curve Analysis System (Growth Curves USA).

### Fluorescence microscopy of CRISPRi strains

Overnight cultures of CRISPRi strains were diluted 1:100 into 200 µL FMC-maltose containing 0.025% xylose, and then incubated at 37°C in a 5% CO_2_ incubator for 16 h. After incubation, 1 µL of 1 mg/mL nile red (ThermoFisher Scientific) was added to each sample and incubated for 5 min at room temperature (RT) in darkness. Next, 1 µL of 1 mg/mL DAPI (4′,6-diamidino-2-phenylindole; ThermoFisher Scientific) was added to each sample and incubated at RT in darkness for a further 5 min. Samples were centrifuged for 5 min at 4,000 x *g* and suspended in ProLong™ Live Antifade Reagent (ThermoFisher Scientific). For imaging, 2.5 µL of stained cells were placed on PBS (pH 7.4; 137 mM NaCl, 2.7 mM KCl, 8 mM Na_2_HPO_4_, 2 mM KH_2_PO_4_) agarose pads [62] before sealing with a coverslip. Images were captured with a Zeiss Axiovert 200M microscope equipped with a Zeiss AxioCam MRm camera using a 100X oil objective.

### Transmission electron microscopy

CRISPRi strains were grown as described for fluorescence microscopy. Afterwards cells were rinsed with 0.1M sodium cacodylate buffer and then fixed in 3% glutaraldehyde overnight at 4°C. The following day, cells were treated with 1.5% osmium tetroxide in the dark for 1 h at 4°C. Next the cells were mixed in equal parts with 5% agarose in PBS, collected by centrifugation at 2000 x *g*, and cooled to 4^°^C. Small chunks of the bacterial pellet plus agarose were then dehydrated in ethanol via the following steps: 30%, 50%, 70%, 80%, 90% each for 15□min, 99% 10 min, and then absolute ethanol 2 × 10 min. The dehydrated cells were embedded in an epoxy resin, sectioned and stained with uranyl acetate and lead citrate. Microscopy was conducted using a Hitachi H7600 transmission electron microscope.

### Waxworm experiments

*Galleria mellonella* killing assays were performed as described previously [63]. Insects in the final instar larval stage were purchased from Vanderhorst, Inc. (St. Marys, OH), stored at 4°C in the dark and used within 7 days of shipment. Groups of 15 larvae, ranging in weight from 200 to 300 mg and with no signs of melanization, were selected at random for subsequent infection. Insects were injected with 1×10^7^ CFU of bacterial inoculum, prepared from overnight cultures that had been washed and resuspended in 0.9% saline solution, into the hemocoel of each larva via the last left proleg using a 25-μl Hamilton syringe. Fifteen larvae were injected for each condition. The number of viable cells injected into the larvae was verified by colony counts on BHI plates. As a negative control, one group of *G. mellonella* were injected with UA159 that was heat killed at 80°C for 30 minutes. After injection, insects were kept at 37°C and survival at different time intervals was recorded. Larvae were scored as dead when thigmotaxis was absent. Kaplan-Meier killing curves were plotted, and estimations of differences in survival were compared using a log rank test. All data were analyzed with GraphPad Prism, version 7.0c software. Experiments were repeated three times.

## Data availability

Raw data for figures, supporting figures and Tn-seq experiments are available in Tables S5-S7.

## Acknowledgements

We thank Dr. Sharon Matthews and Chao Chen at the Electron Microscopy core at the University of Florida for help with TEM imaging and analysis; Dr. Vincent Richards for sending us the *S. mutans* genomes; Dr. Justin Merritt for providing us with the pZX plasmids for xylose-inducible gene expression in *S. mutans*; and Dr. Edward Chan and John Calise for use of, and technical assistance with, fluorescence microscopy. This work was supported by National Institutes of Health grant R01DE013239 (to R.A.B.).

