## Supplemental Text for "Repurposing the *Streptococcus mutans* CRISPR-Cas9 System to Understand Essential Gene Function"

**This PDF file includes:**

Supplemental Materials and Methods

Figs. S1-S15

**Supplemental Materials and Methods**

**Strain cloning.** Site-directed mutagenesis, using the splice overlap extension method [1], was employed to mutate aspartic acid residue 10 to alanine and histidine residue 840 to alanine within *S. mutans cas9* (SMu.1405c). For each mutation, two PCR products were generated using primers that contained the desired mutations, and primers which annealed 0.6 kbp upstream or downstream of the region of interest. The two PCR products, with 25-bp homology that included the mutation, were subjected to PCR for 1 cycle in the absence of added primers. Next, the outer primers were added and a second PCR of 30 cycles was used to amplify the 1.2 kbp fragment with the desired mutation. Both PCR products (for each mutation) were purified and transformed into competent *S. mutans*. Colonies were selected on the basis of erythromycin resistance using a suicide plasmid carrying an internal fragment of the *lacG* gene. Potential clones containing the desired mutations were screened using PCR and DNA Sanger sequencing. Afterwards, strains that had been selected using the *lacG* plasmid were grown in the presence of lactose, to identify strains that had lost the suicide plasmid and the sensitivity of the strains to erythromycin was used as further confirmation of the plasmid. A ∆*cas9* (SMu.1405c) strain was generated in a similar manner, except here the ATG start codon was mutated to a stop codon (TAG; AT>TA). For marking *S. mutans* with GFP, a fragment of DNA was synthesized that contained the P23 promoter and a synthetic RBS followed by a superfolder *gfp* sequence [2]. This fragment was subsequently cloned into the pBGE plasmid that integrates at the *gtfA* locus of *S. mutans* [3]. The restriction enzymes *Xba*I and *Bsr*GI were used to clone the DNA fragment into the pBGE plasmid.

**Green fluorescent protein PAM comparison assay.** PAM sequences were introduced into a P23 promoter driving *gfp* gene expression (pBGE::P23-*gfp*) using the Q5^®^ Site-Directed Mutagenesis Kit. An sgRNA with complementarity adjacent to this PAM was introduced into the pPM::sgRNA plasmid also using the Q5^®^ Site-Directed Mutagenesis Kit. pBGE::P23-*gfp* plasmids (containing PAMs) were introduced into an *S. mutans* strain with a ∆*cas9* mutation and carrying pPM::sgRNA-P23 and P*_xyl_*-d*cas9*_Smu_. GFP fluorescence and microbial growth were measured with a Synergy HT, as detailed in the Materials and Methods section.

**Western blotting.** Bacterial strains were grown in FMC-maltose (with or without xylose) at 37°C to an OD_600_ of 0.5. Cells were harvested by centrifugation at 3,500 × *g* for 10 min, spent medium was discarded, and cell pellets were stored overnight at −20°C. On the following day, the pellets were resuspended in 100 µl of lysis buffer (60 mM Tris, pH 6.8, 2% sodium dodecyl sulfate [SDS]) and transferred to a screw-cap microcentrifuge tube that contained 100 µl 0.1-mm ice-cold glass beads. Samples were homogenized in a bead beater for 30 s three times with 5-min intervals on ice. The samples were then centrifuged at 8,000 × *g* for 10 min at 4°C. The supernates, which contained the *S. mutans* proteins, were carefully removed and placed into a fresh 1.5-ml microcentrifuge tube. The protein concentration was determined using a bicinchoninic acid (BCA) assay following the supplier’s protocol (Pierce), with purified bovine serum albumin used as the standard. Ten µg of each lysate was diluted in 4X SDS loading buffer and the mixture was boiled for 10 min. Proteins were separated by SDS-polyacrylamide gel electrophoresis (PAGE) and then transferred to a polyvinylidene difluoride (PVDF) membrane using a Trans-Blot Turbo transfer system (Bio-Rad). Green fluorescent protein was detected using a polyclonal anti-GFP antibody (1:5,000 dilution; Millipore Sigma) and a goat-anti-rabbit immunoglobulin G (IgG) antibody (1:5,000 dilution; SeraCare Life Sciences, USA). dCas9_Spy_ was detected using a mouse monoclonal anti-CRISPR-Cas9 antibody (1:5,000 dilution; abcam #ab191468) and goat-anti-mouse IgG antibody (1:5,000 dilution; SeraCare Life Sciences, USA). Western blot signals were detected using a SuperSignalWest Pico chemiluminescent substrate kit (Thermo Fisher Scientific) and visualized with a FluorChem 8900 imaging system (Alpha Innotech, USA).

**Essential and growth-supporting genes of *S. mutans* UA159**

Essential genes in *S. mutans* UA159 were identified with Tn-seq performed as described before [4,5], with slight modifications to growth conditions. Transposon mutants were selected on BHI agar, whereas previous libraries were initially selected on blood agar. After libraries were created and stored in frozen aliquots, they were grown in BHI or FMC for 7-10 generations; previous libraries were passaged for ~30 generations. The results of these experiments were used to determine which genes to target with CRISPRi in parallel with previous Tn-seq in *S. mutans* UA159. In addition, for poorly annotated essential or growth-supporting genes we ran the protein sequence through BLASTP.

**RNA-sequencing**

RNA-sequencing was performed as described previously by Zeng et al [6]. Briefly, RNA was extracted from OD_600_ bacterial cultures using the RNeasy Mini Kit. Next, RNA was treated with the MICROB*Express* Bacterial mRNA Enrichment Kit (ThermoFisher Scientific) to remove 16S and 23S rRNAs. After this step the quality of mRNA was assessed using an Agilent Bioanalzyer (Agilent Technologies) at the University of Florida NGS core facility. Following the quality check, cDNA libraries were made using the NEBNext Ultra II Directional RNA Library Prep Kit and NEBNext Multiplex Oligos for Illumina (New England Biolabs). Deep sequencing was performed by the University of Florida NGS core facility on an Illumina NextSeq500 DNA sequencing machine. After sequencing, read counts were aligned to the *S. mutans* UA159 genome using bioinformatics tools hosted on the Galaxy server maintained by the Research Computing Center at the University of Florida. Gene expression changes between samples were quantified with Degust (http://degust.erc.monash.edu/) using the Voom/Limma methodology.

The original RNA-seq data from this study was uploaded to the GEO database (<https://www.ncbi.nlm.nih.gov/geo/>) with the accession number GSE141863.

**Bacterial two-hybrid assays**

Bacterial adenylate cyclase two-hybrid (BACTH) assays were performed by following supplier’s instructions (Euromedex). A full description of this bacterial two-hybrid system can be found in the paper by Karimova et al [7]. Briefly, genes of interest were cloned into the two adenylate cyclase plasmids pUT18C and pKT25 (see S4 Table for primer sequences). This cloning was performed in *E. coli* 10-beta. After cloning was validated with Sanger sequencing of the cloned regions, plasmid constructs were transformed into electrocompetent *E. coli* BTH101. For the transformation, both plasmids pUT18C and pKT25 were co-transformed simultaneously. After selecting for co-transformations (LB plus 100 µg/mL ampicillin and 50 µg/mL kanamycin), colonies were grown overnight in LB media. The following day 5 µl of each interaction strain was plated onto M9 agar containing glucose and 50 µg/mL 5-Bromo-4-chloro-3-indolyl-β-D-galactopyranoside (X-gal). Agar plates were incubated at 30°C for 16 h overnight and imaged. Transformations were done in triplicate with selected images representative of three biological replicates.

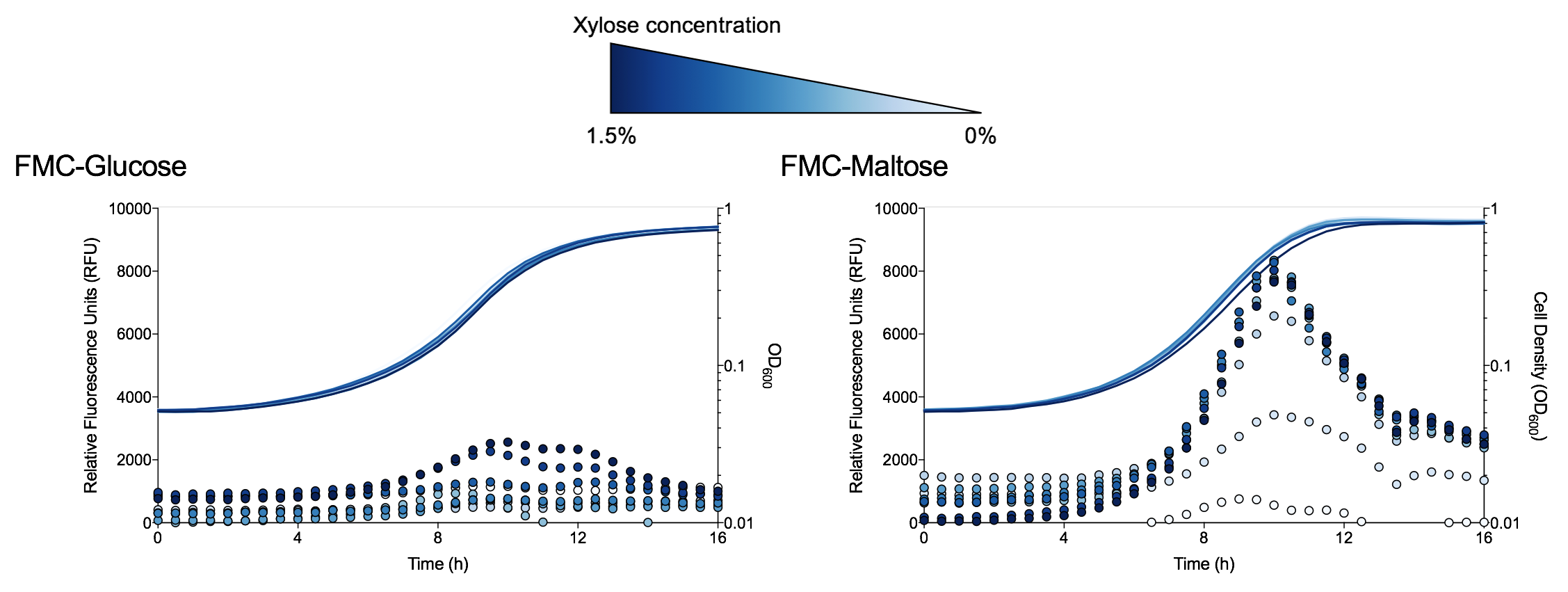

**Figure S1.** **Determining the optimal conditions for xylose induced gene expression in *S. mutans*.** The graphs show (glucose left, maltose right) that P*_xyl_*-*gfp* expression was higher in FMC-maltose than FMC-glucose for comparable amounts of xylose. Fluorescence readings are shown as dots, and cell density readings as lines. As the xylose concentration is diluted data points change from a dark to a light blue color.

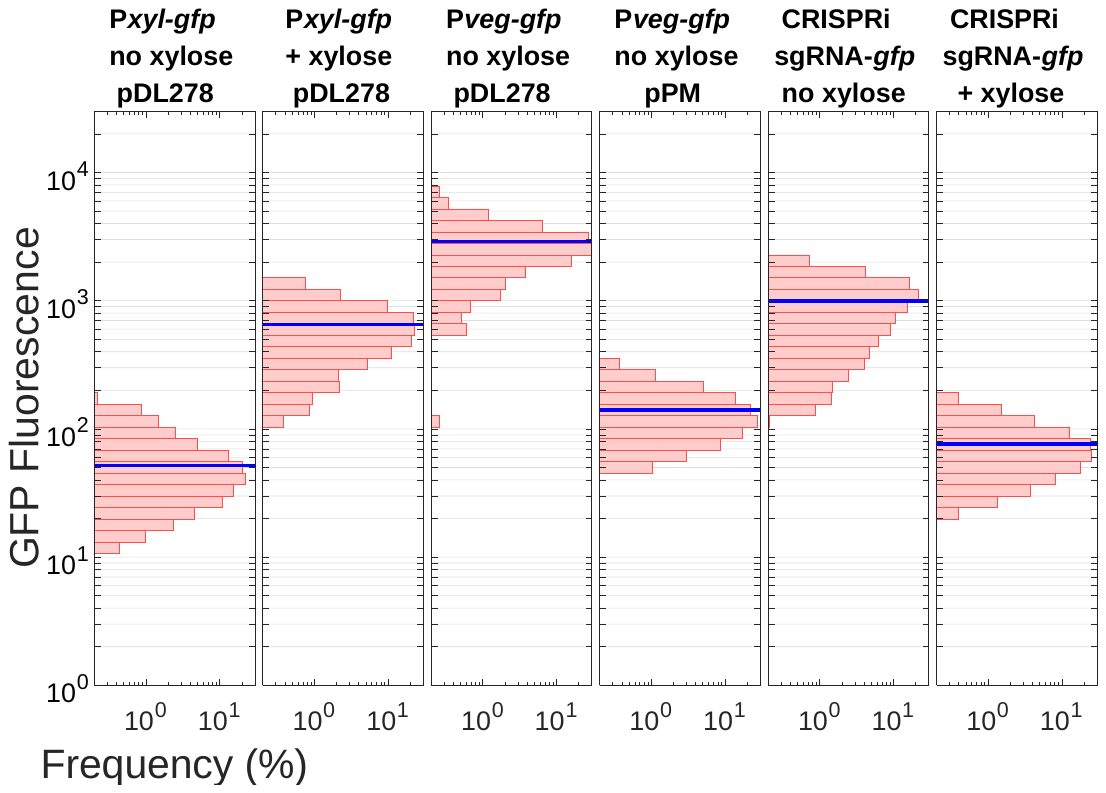

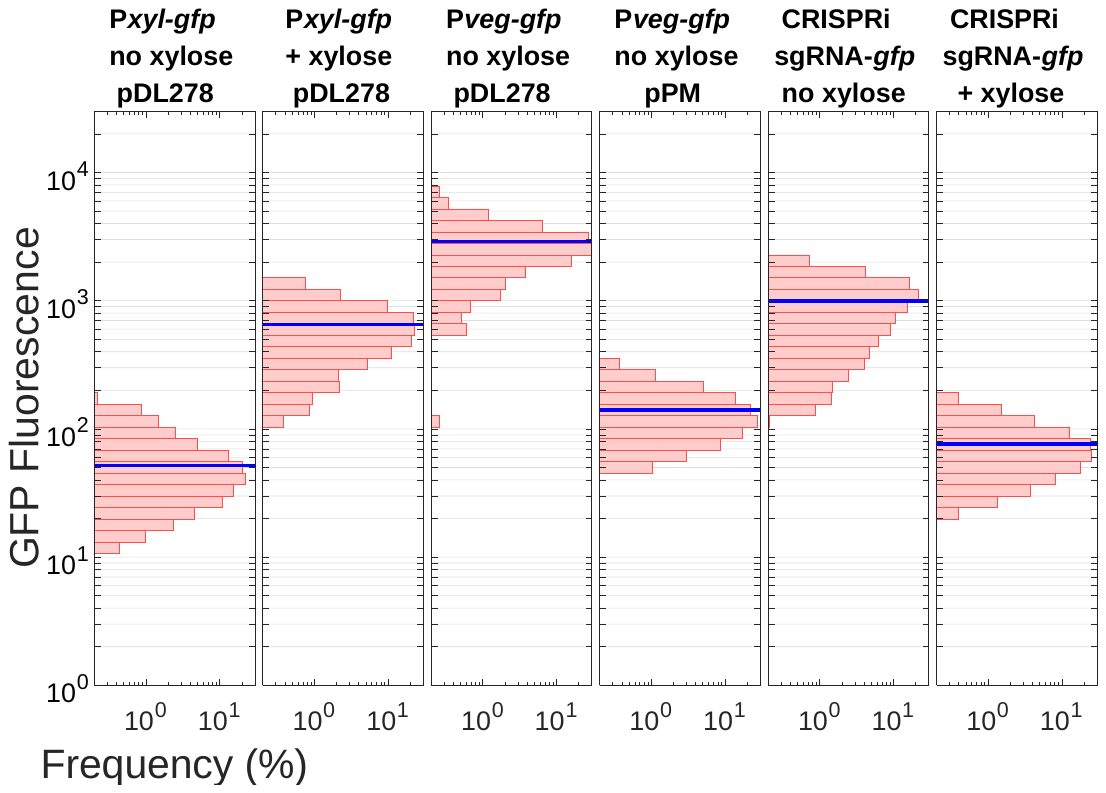

**Figure S2. Single-cell analysis of P*_xyl_*-*gfp* expression and CRISPRi knockdown.** Addition of xylose to FMC-maltose induced *gfp* expression in virtually all imaged cells (first versus second panel) although *gfp* expression was not as high as observed with a constitutive P*_veg_* promoter (third panel). In a CRISPRi strain containing a chromosomally integrated *gfp* reporter all cells were GFP positive without xylose addition (fourth panel). Virtually no cells expressed *gfp* when xylose was added to the CRISPRi strain as median GFP fluorescence was barely above background (fifth panel).

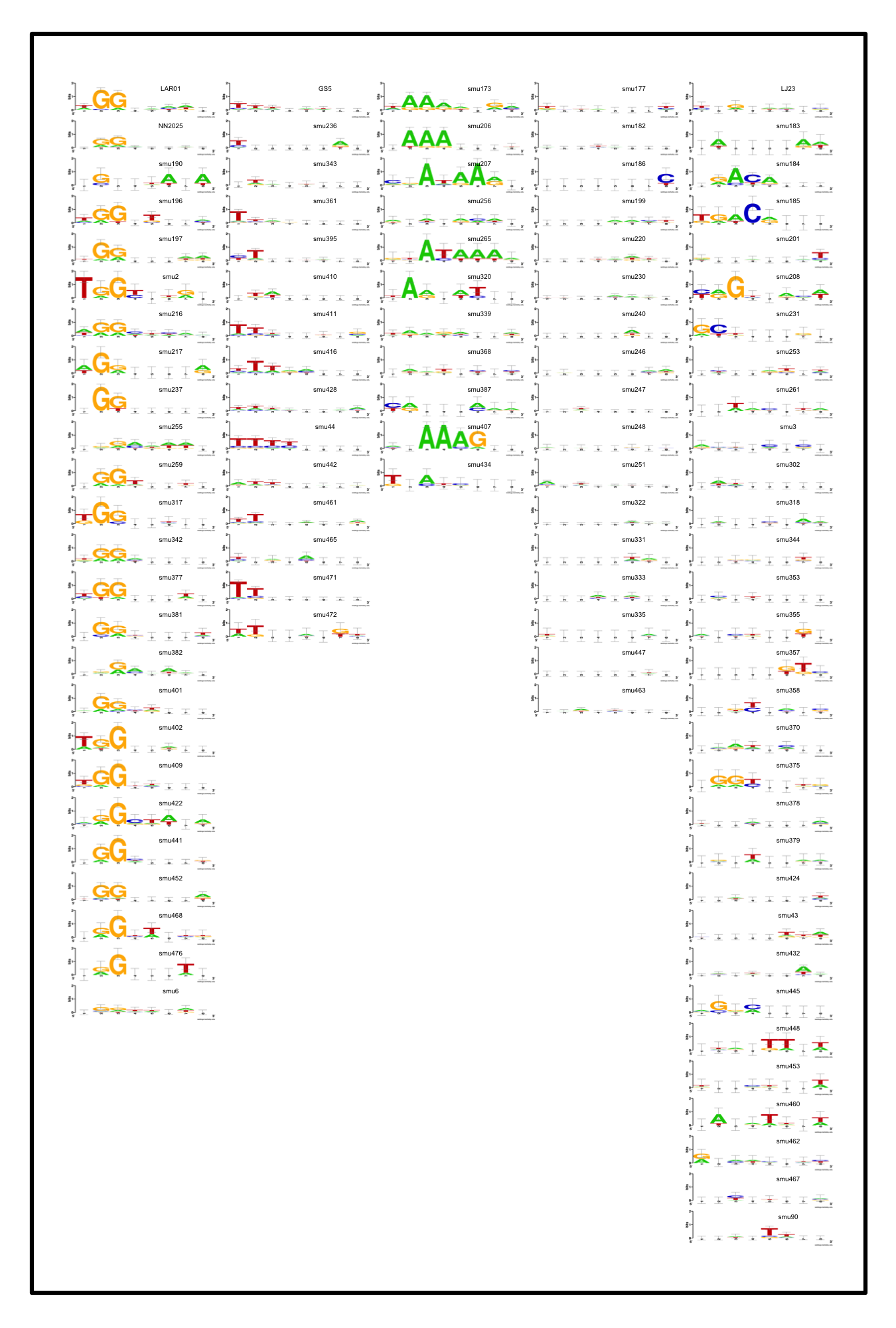

**Figure S3. Cas9 PAM predictions for *Streptococcus mutans* strains.** Spacer sequences found within Type II CRISPR cassettes of *S. mutans* strains were aligned to the genome of the *Streptococcus* phage M102AD. After alignment protospacer adjacent motifs (PAMs) were extracted and consensus PAMs were generated with WebLogo. Consensus PAMs broadly fit into five groups, 5’-NGG-3’, T-rich, A-rich, no distinct PAM and a fifth group of PAMs that didn’t fit into the first four groups.

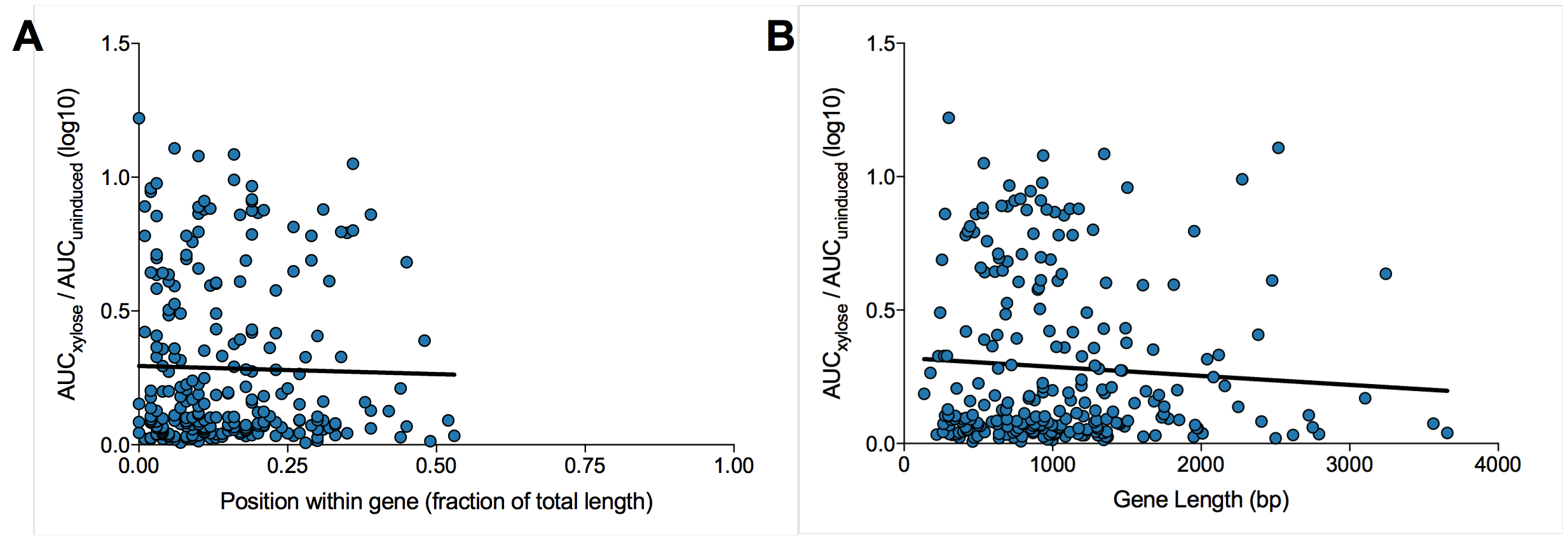

**Figure S4. Investigating the impact of sgRNA position within gene and overall gene length on CRISPRi growth defects.** (A) Overall, there was no discernable impact on CRISPRi mediated growth defects according to the placement of the sgRNA within the gene. (B) Gene length did not impact CRISPRi-mediated growth defects across all measured strains (*e.g.* larger genes showing stronger growth defects versus smaller genes).

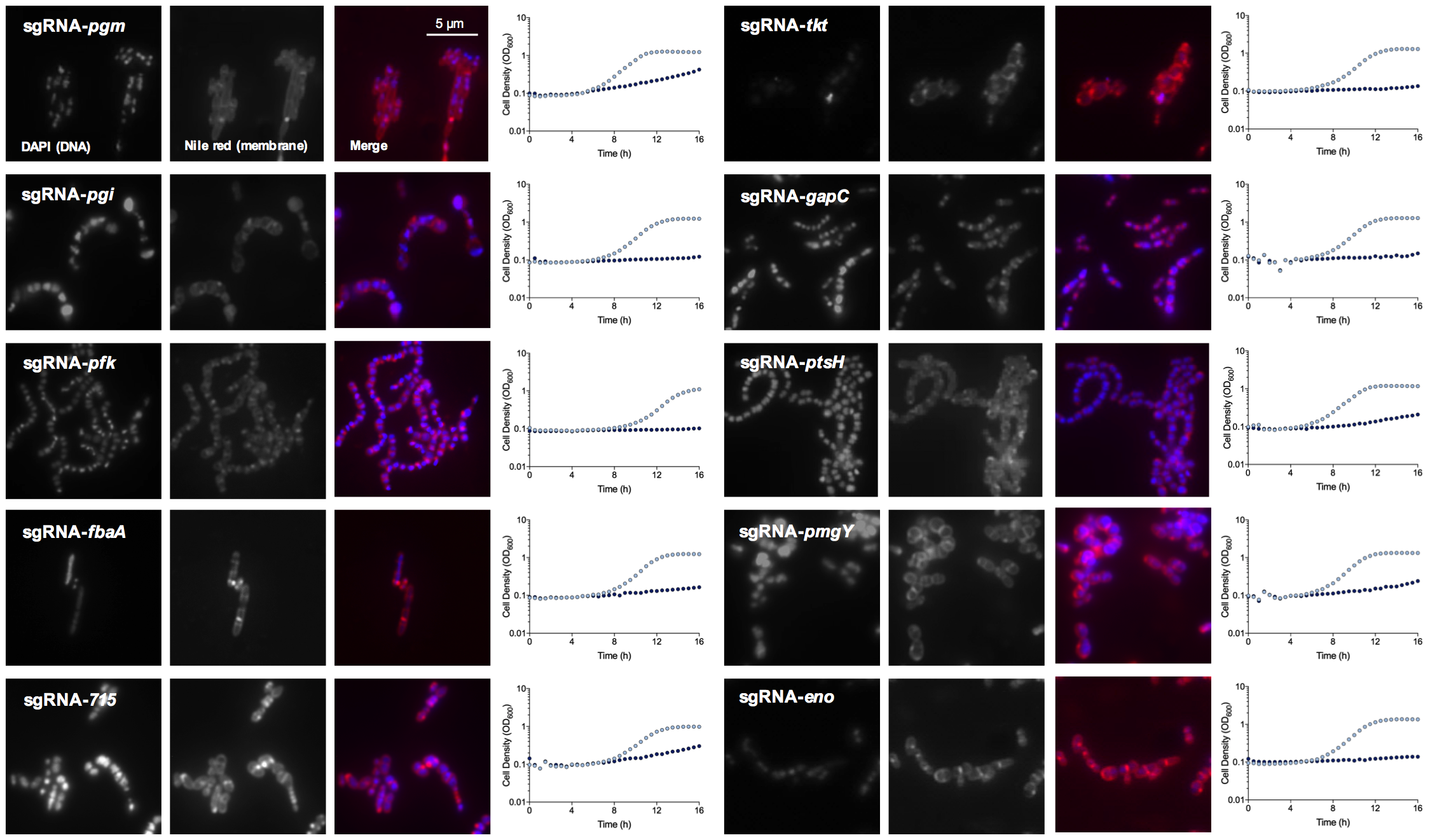

**Figure S5. Growth phenotypes and microscopy of CRISPRi strains with silenced genes related to carbohydrate metabolism.** For growth assays, light blue dots indicate the no xylose condition and dark blue dots represent the addition of 0.1% xylose. Cells were stained with DAPI (DNA stain) and nile red (membrane stain) and imaged with a x100 oil objective. For each strain a representative microscopic image is shown.

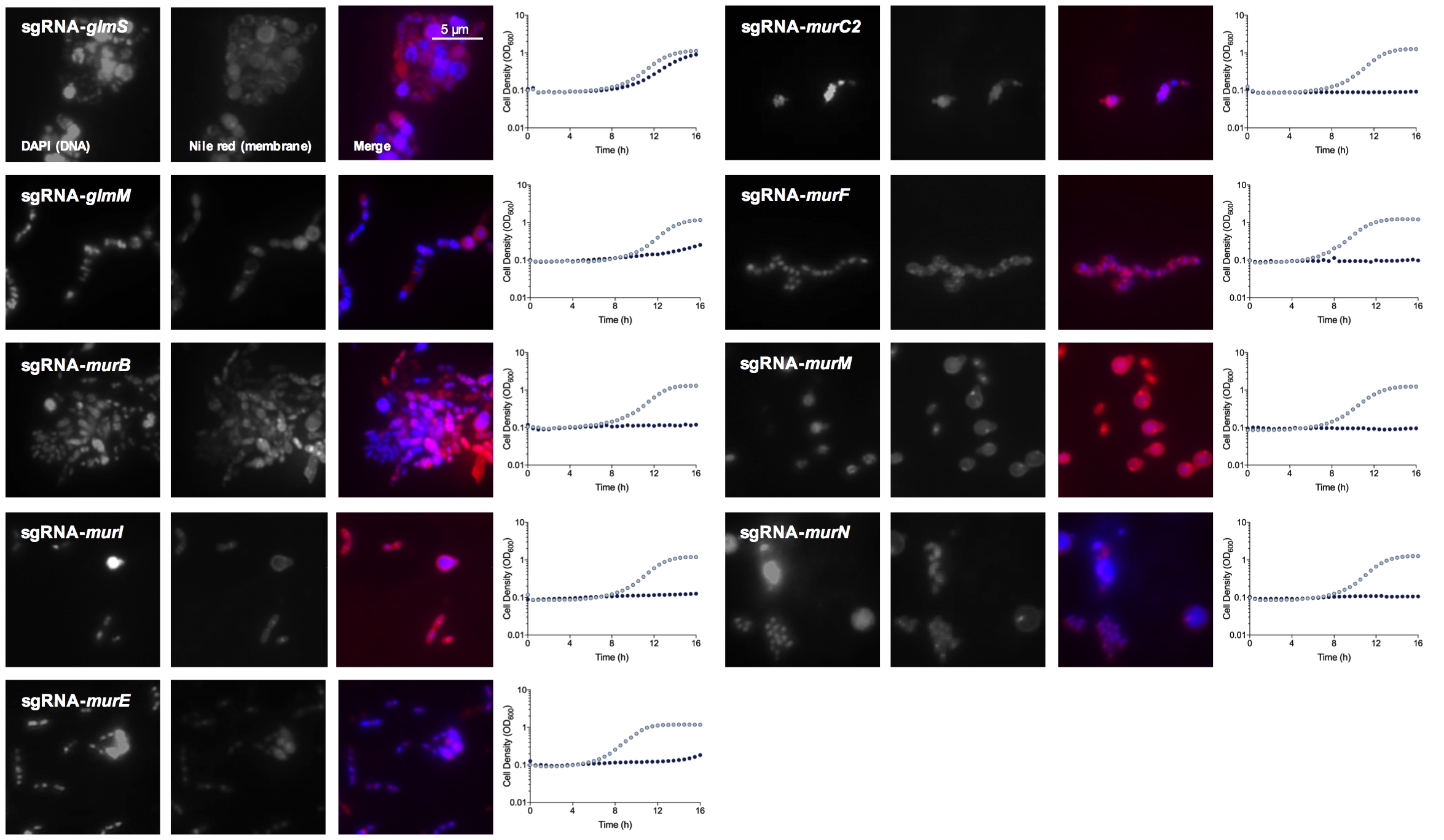

**Figure S6. Growth phenotypes and microscopy of CRISPRi strains with silenced genes related to peptidoglycan biosynthesis.** For growth assays, light blue dots indicate the no xylose condition and dark blue dots represent the addition of 0.1% xylose. Cells were stained with DAPI (DNA stain) and nile red (membrane stain) and imaged with a x100 oil objective. For each strain a representative microscopic image is shown.

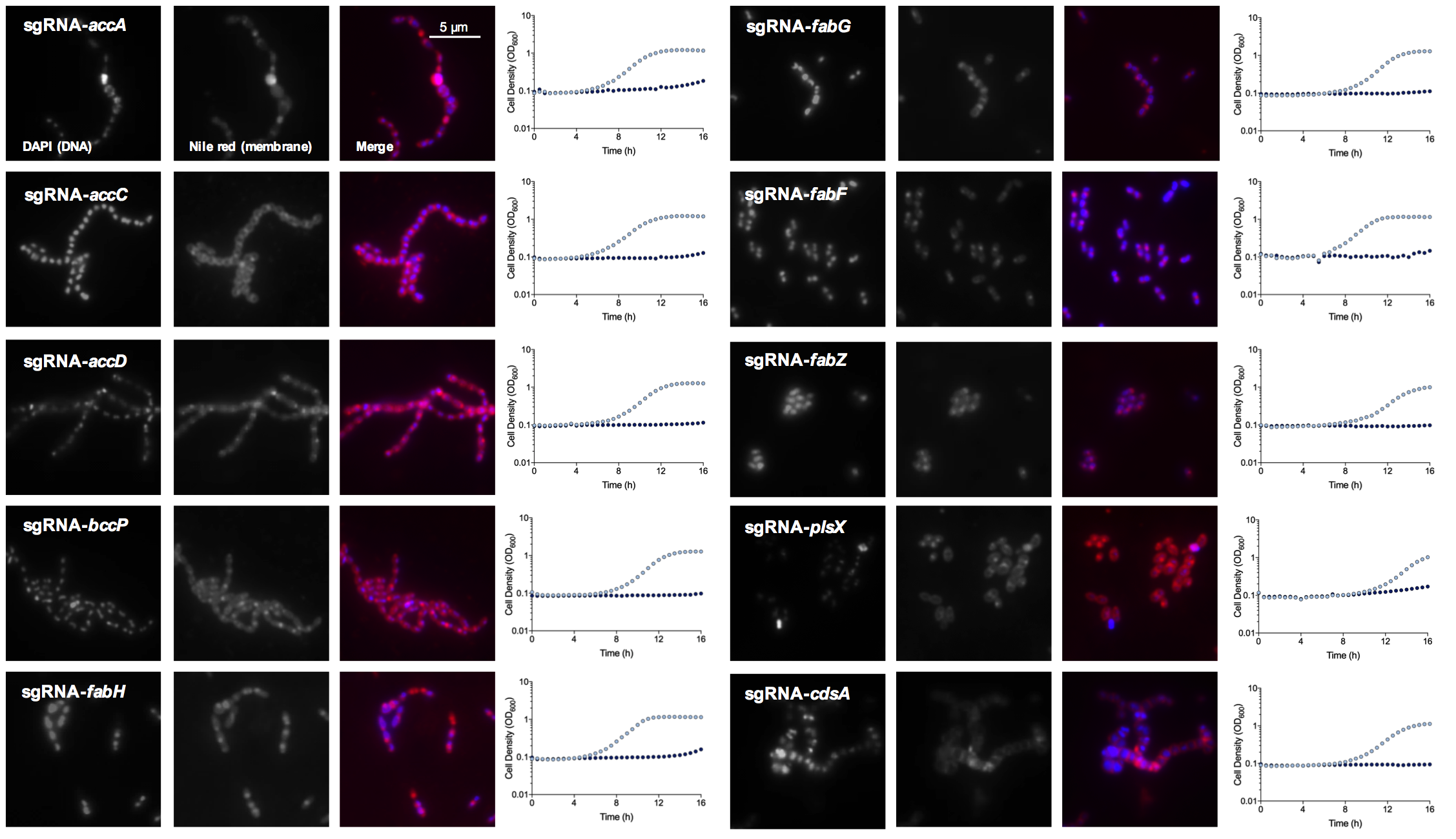

**Figure S7. Growth phenotypes and microscopy of CRISPRi strains with silenced genes related to lipid metabolism.** For growth assays, light blue dots indicate the no xylose condition and dark blue dots represent the addition of 0.1% xylose. Cells were stained with DAPI (DNA stain) and nile red (membrane stain) and imaged with a x100 oil objective. For each strain a representative microscopic image is shown.

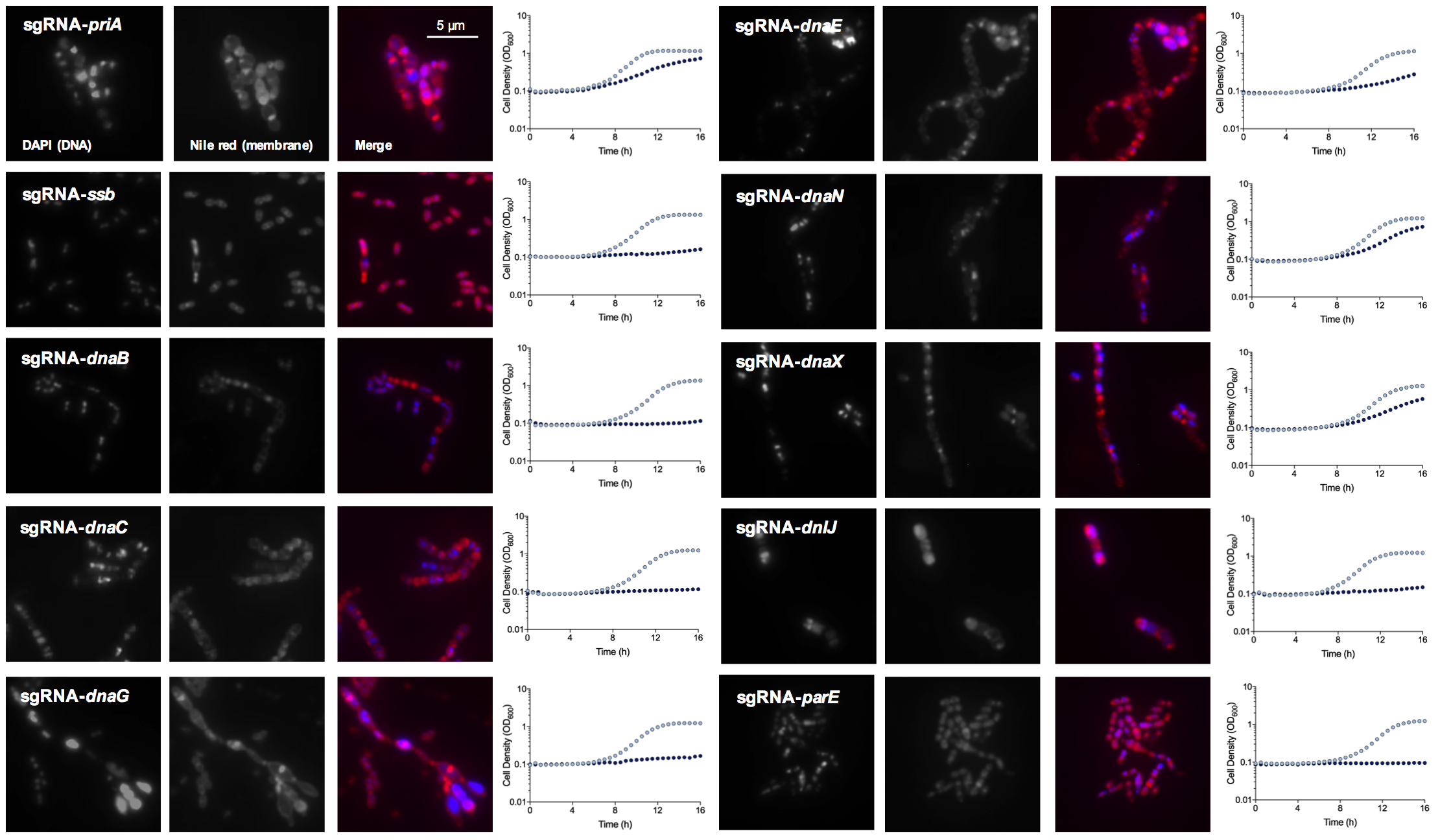

**Figure S8. Growth phenotypes and microscopy of CRISPRi strains with silenced genes related to DNA replication.** For growth assays, light blue dots indicate the no xylose condition and dark blue dots represent the addition of 0.1% xylose. Cells were stained with DAPI (DNA stain) and nile red (membrane stain) and imaged with a x100 oil objective. For each strain a representative microscopic image is shown.

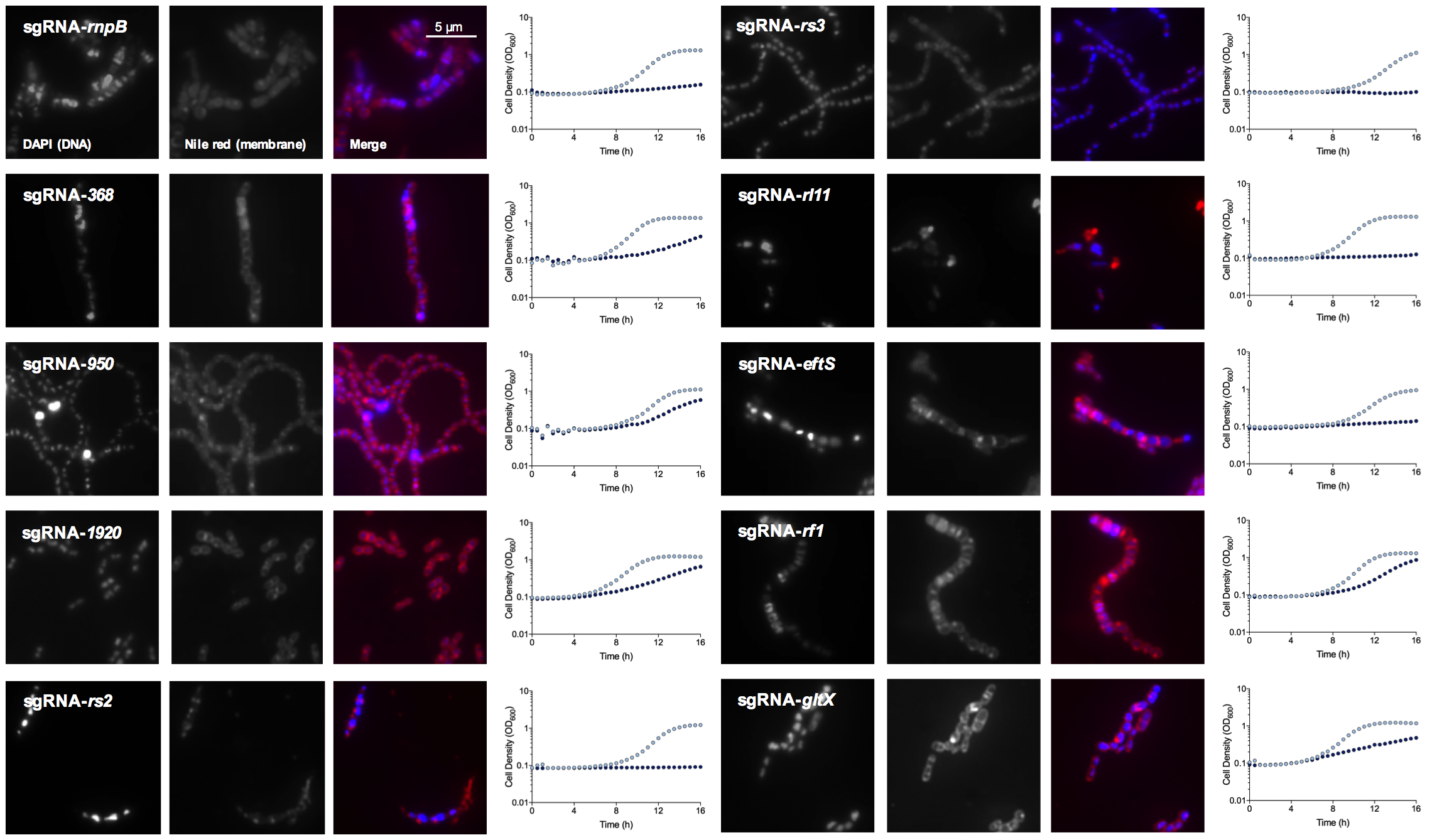

**Figure S9. Growth phenotypes and microscopy of CRISPRi strains with silenced genes related to translation and ribosome biogenesis.** For growth assays, light blue dots indicate the no xylose condition and dark blue dots represent the addition of 0.1% xylose. Cells were stained with DAPI (DNA stain) and nile red (membrane stain) and imaged with a x100 oil objective. For each strain a representative microscopic image is shown.

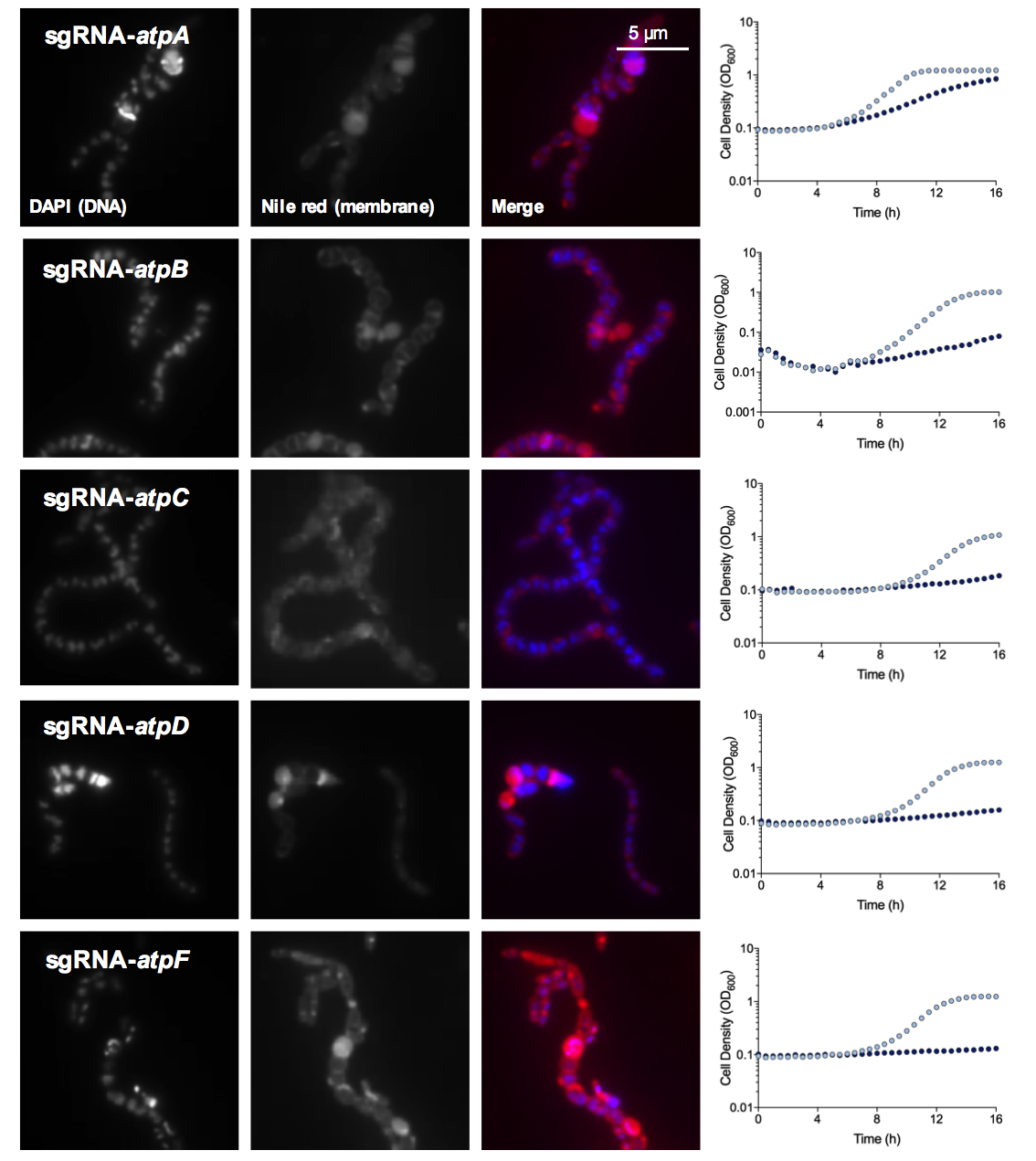

**Figure S10. Growth phenotypes and microscopy of CRISPRi strains with silenced genes related to ATP synthase activity.** For growth assays, light blue dots indicate the no xylose condition and dark blue dots represent the addition of 0.1% xylose. Cells were stained with DAPI (DNA stain) and nile red (membrane stain) and imaged with a x100 oil objective. For each strain a representative microscopic image is shown.

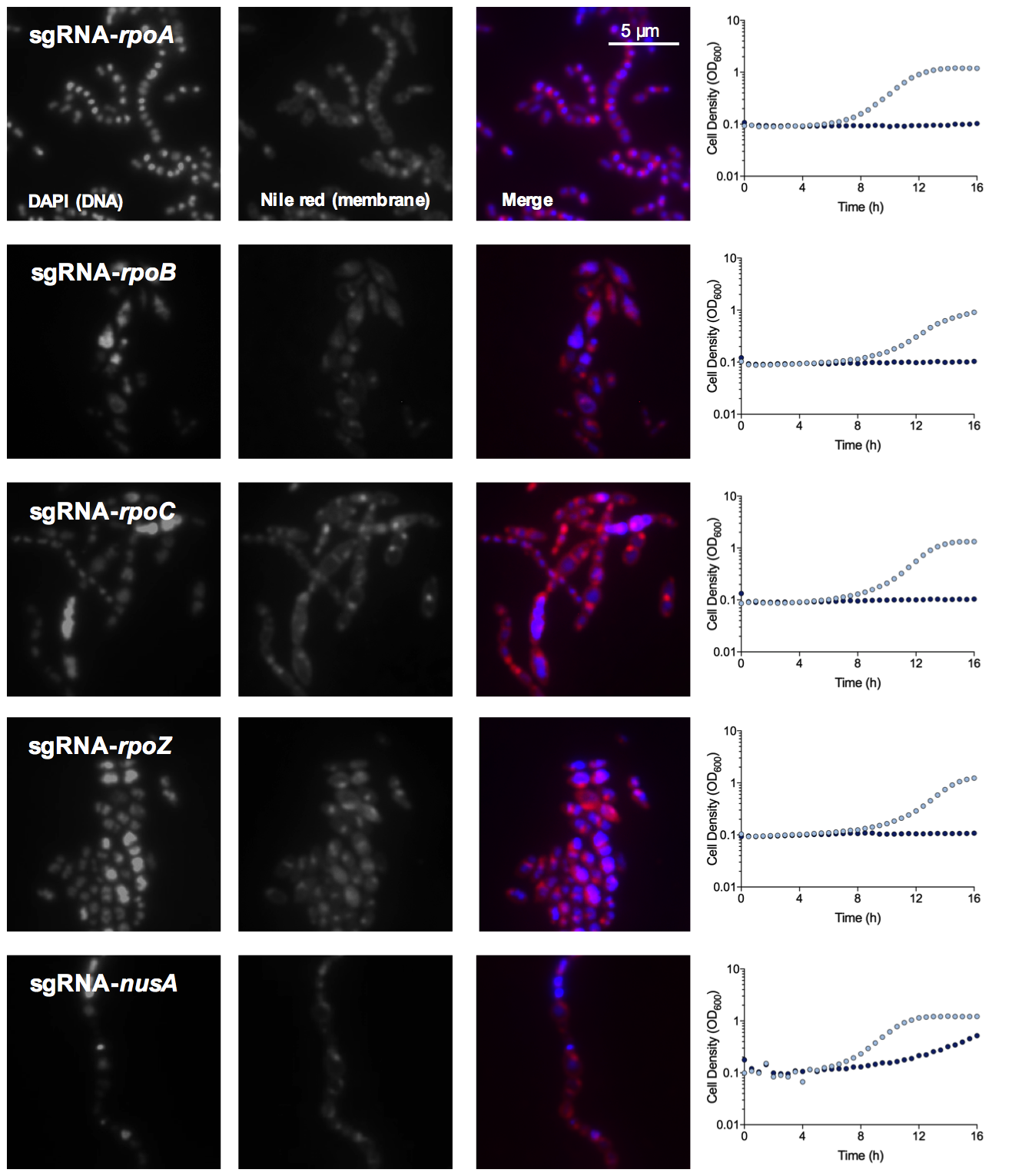

**Figure S11. Growth phenotypes and microscopy of CRISPRi strains with silenced genes related to transcription.** For growth assays, light blue dots indicate the no xylose condition and dark blue dots represent the addition of 0.1% xylose. Cells were stained with DAPI (DNA stain) and nile red (membrane stain) and imaged with a x100 oil objective. For each strain a representative microscopic image is shown.

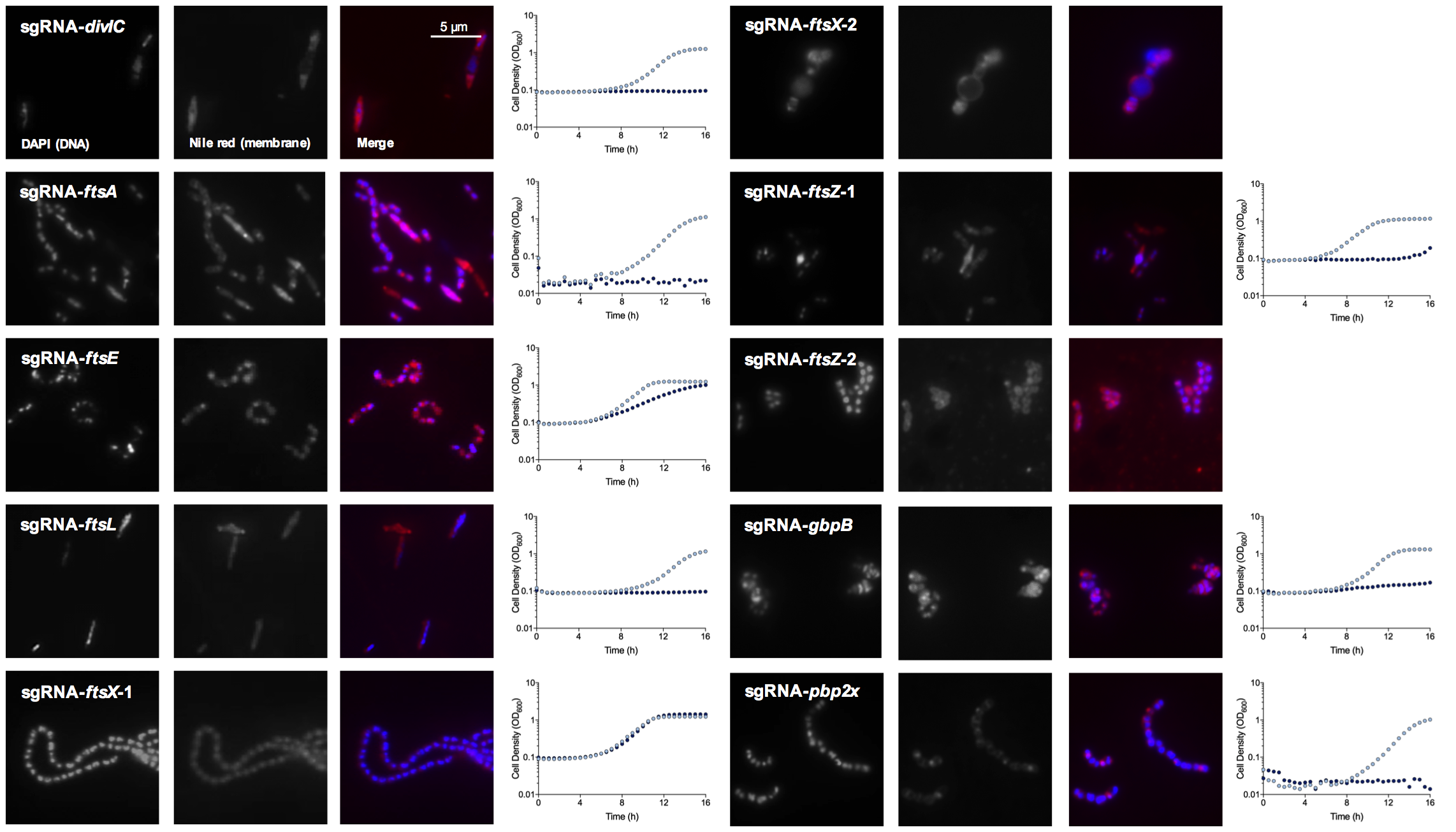

**Figure S12. Growth phenotypes and microscopy of CRISPRi strains with silenced genes related to cell division.** For growth assays, light blue dots indicate the no xylose condition and dark blue dots represent the addition of 0.1% xylose. Cells were stained with DAPI (DNA stain) and nile red (membrane stain) and imaged with a x100 oil objective. For each strain a representative microscopic image is shown.

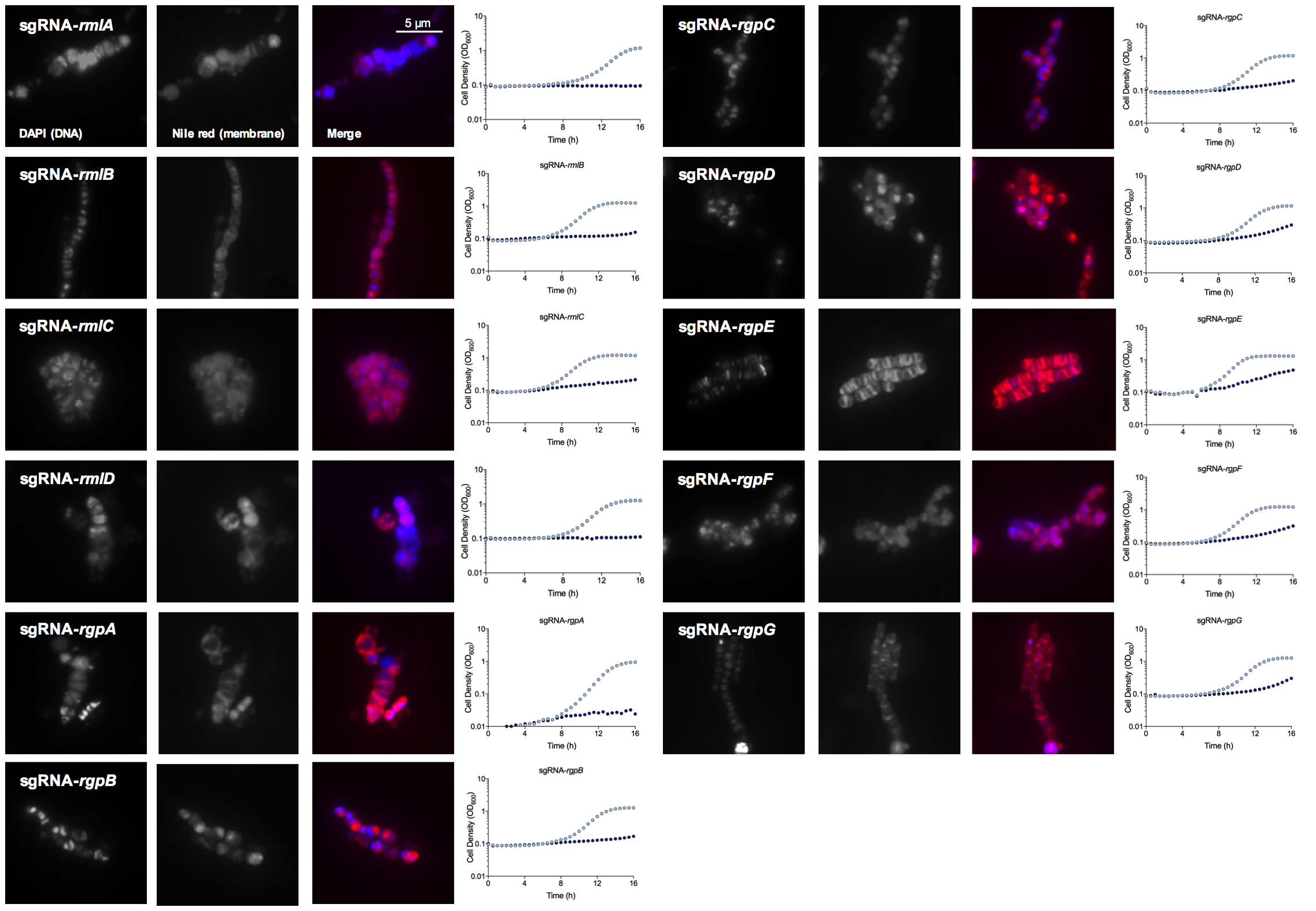

**Figure S13. Growth phenotypes and microscopy of CRISPRi strains with silenced genes related to rhamnose-glucose polysaccharide biosynthesis and transport.** For growth assays, light blue dots indicate the no xylose condition and dark blue dots represent the addition of 0.1% xylose. Cells were stained with DAPI (DNA stain) and nile red (membrane stain) and imaged with a x100 oil objective. For each strain a representative microscopic image is shown.

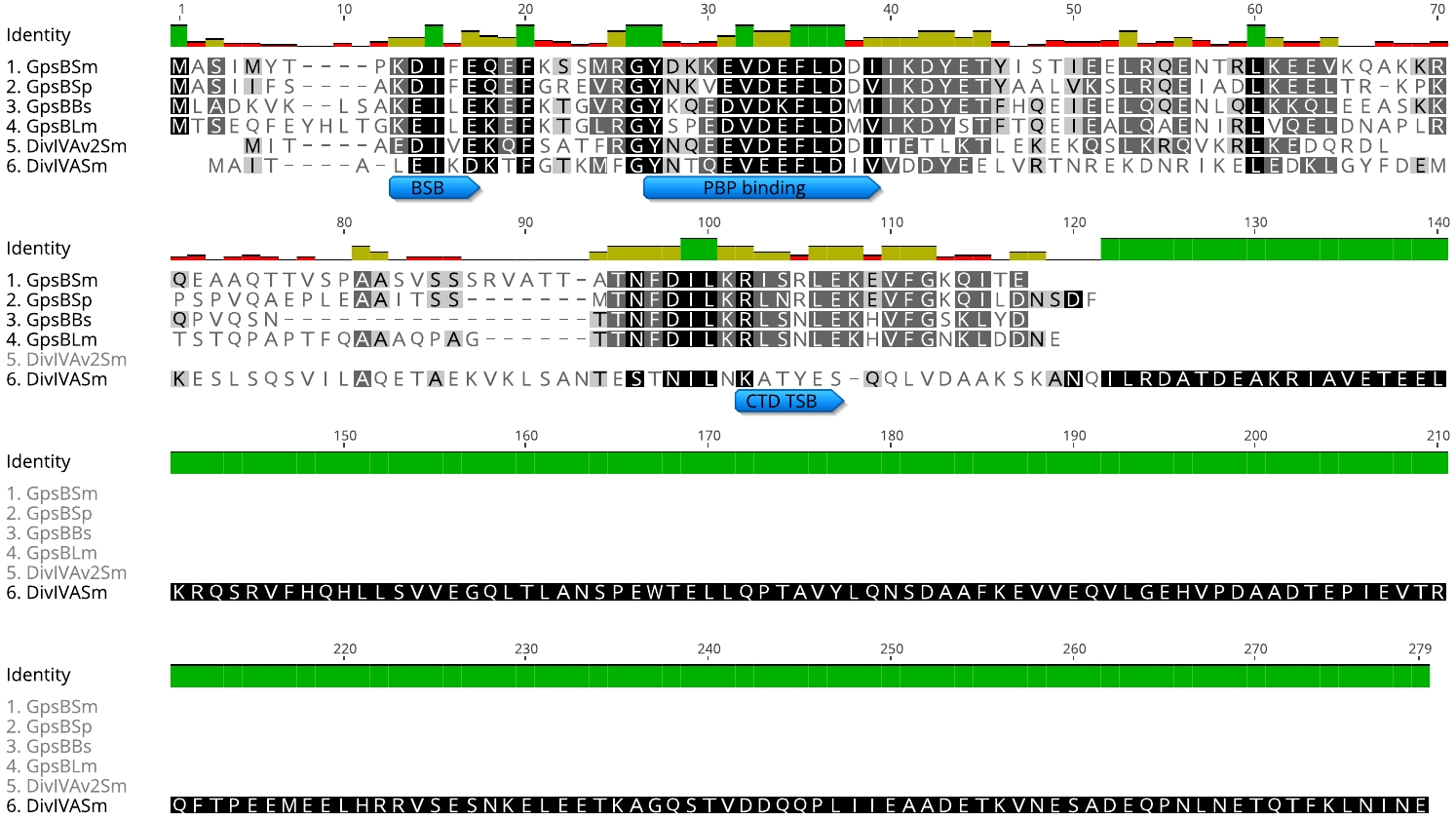

**Figure S14. Alignment of SMu.471 protein sequence with Gram positive GpsB proteins.** (1) *Sm*, *Streptococcus mutans* (GpsB, SMu.471); (2) *Sp*, *Streptococcus pneumoniae*; (3) *Bs*, *Bacillus subtilis*; (4) *Lm*, *Listeria monocytogenes*. Regions with increased conservation are shown with darker grey/black highlighting. Conserved domains are shown: GpsB bidentate salt bridge (BSB), penicillin binding protein binding (PBP binding), and C-terminal domain trimer salt bridge (CTD TSB). The protein sequence of SMu.471 was also compared to two additional *S. mutans* proteins that contain DivIVA domains, DivIVAv2Sm (5; SMu.1260c), and DivIVASm (6; SMu.557).

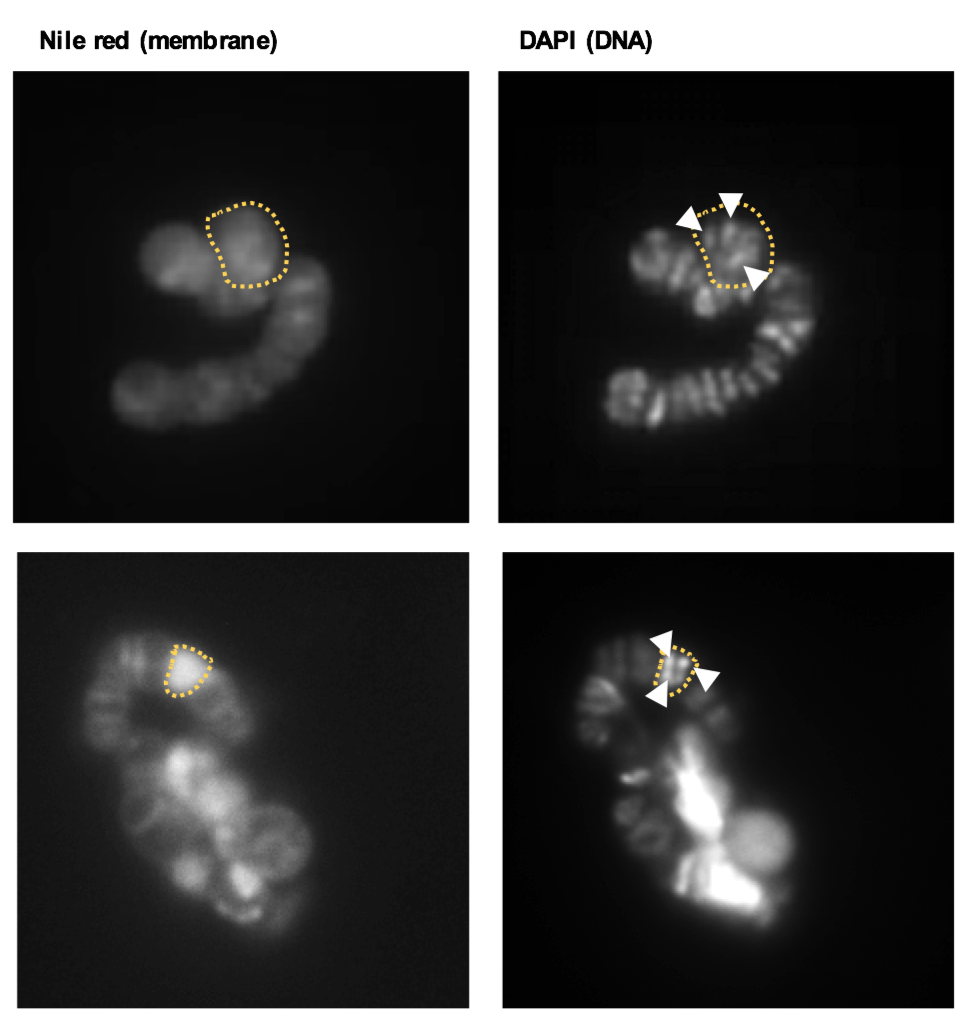

**Figure S15. Multiple nucleoids per cell for RGP biosynthesis knockdowns.** Cell borders are shown with dotted orange lines. Inside outlined cells multiple nucleoids are visible as punctate bright dots, also highlighted with white arrowheads.

**Other Supporting Information Files**

Table S1 Transcriptomic changes measured by RNA-seq in CRISPRi strains

Table S2 Database of essential genes and CRISPRi growth phenotypes

Table S3 Strains and plasmids used in this study

Table S4 Oligonucleotides used in this study

Table S5 Raw data for figures

Table S6 Raw data for supporting figures

Table S7 Number of transposon reads per gene
